# WASP: A software package for correctly characterizing the topological development of ribbon structures

**DOI:** 10.1101/2020.09.17.301309

**Authors:** Zachary Sierzega, Jeff Wereszczynski, Chris Prior

## Abstract

We introduce the Writhe Application Software Package (WASP) which can be used to characterise the topology of ribbon structures, the underlying mathematical model of DNA, Biopolymers, superfluid vorticies, elastic ropes and magnetic flux ropes. This characterisation is achieved by the general twist-writhe decomposition of both open and closed ribbons, in particular through a quantity termed the polar writhe. We demonstrate how this decomposition is far more natural and straightforward than artificial closure methods commonly utilized in DNA modelling. In particular, we demonstrate how the decomposition of the polar writhe in local and non-local components distinctly characterizes local helical structure and knotting/linking of the ribbon. This decomposition provides additional information not given by alternative approaches. As an example application, the WASP routines are used to characterise the evolving topology (writhe) of DNA minicircle and open ended plectoneme formation magnetic/optical tweezer simulations. Finally it is demonstrated that a number of well known alternative writhe expressions are actually simplifications of the polar writhe measure.

## Introduction

Quantifying the varying and complex geometries of three dimensional curves is an important task in many fields. For example flexible biomolecular structures, such as polymer chains, DNA helices, and chromatin fibers, adopt a wide range of conformations when exposed to different solvent/cosolute environments and external forces and torques^1–3^. Characterizing the evolving morphology of these chains poses significant mathematical challenges that naturally draw on the fields of topology and differential geometry.

One of the most useful metrics for quantifying large scale structural changes in these complexes is writhe. The writhe is a global geometric quantity commonly used to characterize the conformational variety of circular DNA structures^4^. Its utility relates to the fact that, for ribbon structures (see Figure 1(a)) which can be used to model structures from DNA molecules to magnetic flux ropes, it forms part of the invariant sum:

**Figure 1.**
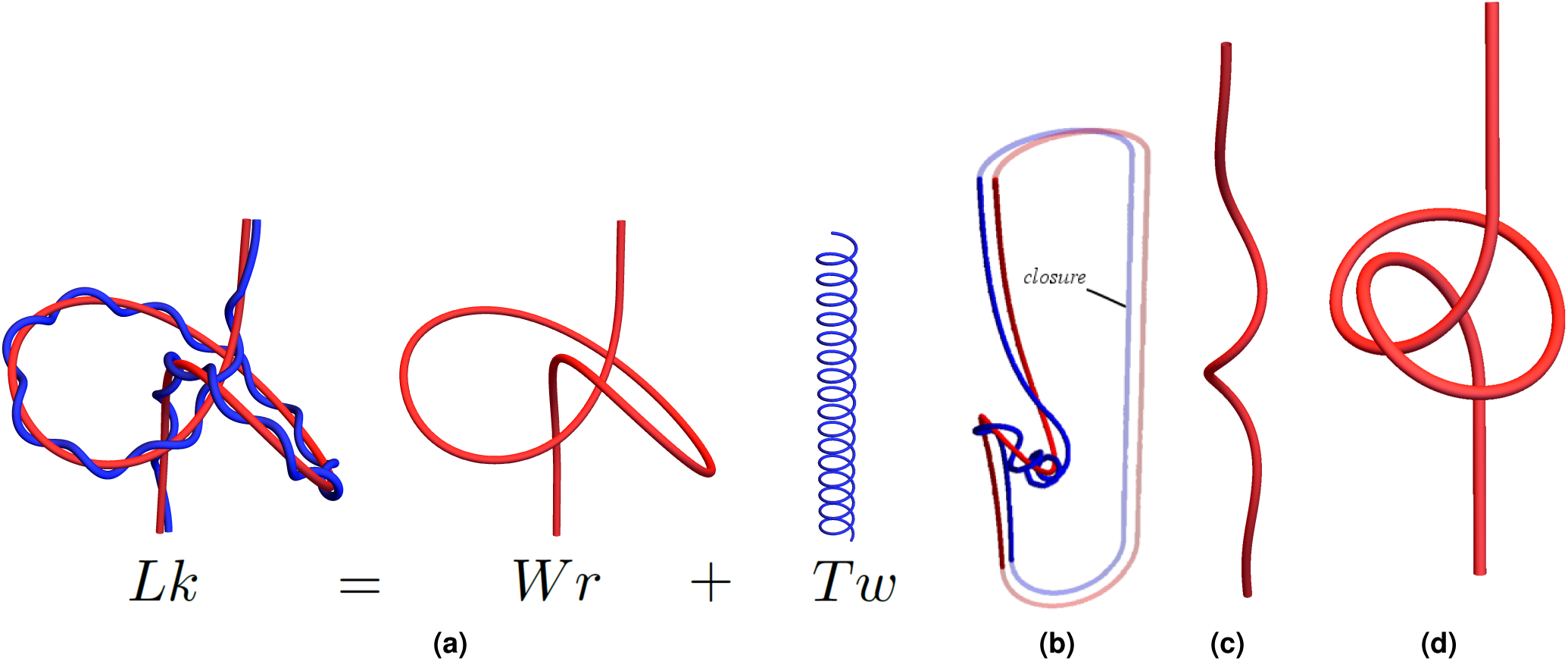
Illustrations of concepts discussed in the introduction. Panel (a), is an illustration of the meaning of the *Wr* + *Tw* decomposition. The left figure is a ribbon a structure composed of an axis curve (red) and a second curve wrapping around this axis (Blue). The linking of these two curves (invariant if the ends of the ribbon are fixed) can be decomposed into the self linking of the ribbon’s axis (*Wr*) and the total rotation of the second curve around the axial direction of the first *Tw*. Panel (b) indicates an artificial extension of the ribbon (a closure). Panels (c) and (d) illustrate what is meant by local and non-local writhing. The curve in panel (c) coils helically at its centre, the local writhe measures this helical coiling along the curve’s length. The curve in panel (d) is knotted/self entangled, that is to say distinct sections of the curve wrap around each other. This is non-local writhing.

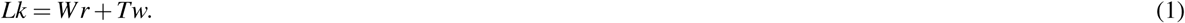

Here *Lk* is the linking number, a topological measure of the entwined nature of the two edges of the ribbon which is unchanged under any change in shape which forbid their crossing (*i*.*e* it categorizes the ribbon’s entanglement). The writhe *Wr* represents the contribution to *Lk* from the self linking of the axis curve on itself (Figure 1(a)). The twisting *Tw* is a measure of the rotation of the ribbon about its axis^5^ (Figure 1(a)). As an example of its use, the *Lk* can be used as a fixed model constraint on the number of DNA Sequence repeats. The constraint is then applied to an energy models of the DNA backbone, which then often has the twisting *Tw* as an elastic energy component^6–11^. The writhe can then be used to constrain the allowed global (axial) shape of the molecule. In DNA models writhe measures supercoiling.

The most commonly known version of the link-twist-writhe relationship, equation (1), is for closed ribbon/curves such as DNA mini-circles. it was originally derived by G Călugăreanu^12–14^ and was popularised (but not derived) for a biological audience by F B Fuller^5,15^. However, the theorem as originally derived is not applicable to open-ended ribbons such as those shown in Figure 1(a) as the linking (*Lk*, as defined in that version of the theorem) is not an invariant for open-ended ribbon structures^16^. Therefore, it cannot constrain the interplay between internal twisting (*Tw*) and global self entanglement of the ribbon’s axis (*Wr*). Many target applications which can be modelled by ribbons are open structures, such as DNA molecules subjected to optical tweezer experiments and chromatin fibers.

To overcome this problem, Fuller originally suggested that one could extend the closed ribbon theorem to open ribbons by artificially extending the structure^15^ (as shown in Figure 1(b)). This approach has been popular in the field of DNA modelling^7,8,17–21^ and elastic tube applications^22–24^. It is complicated by the fact that the closure will generally contribute to both the writhing and linking of the composite, and hence the the extraction of clear geometrical insight from the calculated quantities cannot be consistently achieved^16,25,26^. Alternative approaches have been formulated which approximate the writhing value^15,17,23,27,28^, however, these will in general not charcterize the writhing^16,25^ and hence cannot consistently be used to constrain the topology of the ribbon. Finally, some authors have simply chosen to utilize the definitions of the original closed formula without a closure in this case the *Lk* measure is no longer a topological invariant so the relationship looses its fundamental topological anchoring (the sum *Tw* + *Wr* is no longer fixed).

Previously, Berger and Prior^16^ introduced a version of (1) which **is** applicable to open-ended ribbons. It is based on a definition of *Lk* (called the net-winding) which is invariant to all deformations of the ribbon which do not permit rotation at its ends. In supercoiling ribbon models it would be equivalent to the natural linking of the DNA plus the applied over-coiling of the structure. The twisting *Tw* is the same quantity as in the closed case and a definition of the writhing (termed the polar writhe) was based on the difference of these quantities. For closed curves, these quantities have the same value as those of the closed variant of (1) so to this is a more general topological relationship than the original as it contains it but is applicable in a wider range of applications. This framework has been regularly applied in Solar physics applications since the original paper *e*.*g*.^29–35^ and is increasingly being applied in Elastic tube modelling^26,36,37^. However, it has not been widely adopted in the biophysical community, as artificial closures methods are still readily employed for problems such as characterizing DNA supercoiling.

Here, we seek to introduce polar writhe to the biophysical community and to provide a user-friendly tool for its implementation. The first aim of this article is to introduce the open-ribbon version of equation (1) derived in^16^, and extended to consider knotting deformations in^26^. Both papers are somewhat technical, necessarily involving mathematical proofs. In this note we illustrate the properties of the polar writhe which underpin this open decomposition though instructive examples of constrained DNA supercoiling simulations. The second aim is to demonstrate how decomposing the polar writhe into local and non-local components, which measure “spring-like extension” deformations and “buckling-type” deformations/supercoiling changes respectively (see Figure 1(c) and (d)), can provide critical additional insight into the curve’s geometry through a series of instructive examples. Third, we demonstrate how an extension of (1), as defined in^26^ can detect transitions to knotting possible in open ribbon structures, even when its ends are constrained from moving. Finally, we introduce the *Writhe Analysis Software Package* (WASP) which can calculate the components of equation (1), both in the closed and open ended case. WASP allows non-expert users to easily apply these calculations to standard curve files and is particularly geared towards usage with molecular dynamics trajectories, supporting common trajectory file types including those generated by popular MD software including AMBER and OxDNA as well as more general MD file types such as PDB files.

The paper is structured as follows. In the first section, we introduce the polar writhe through a series of instructive example calculations. This includes highlighting the local-non local decomposition and the additional information it provides. The Methods section provides a description of the DNA simulations experiments we performed, which were designed to highlight aspects of the open *Lk*-*Tw*-*Wr* decomposition described in^16,26^. This includes a description of how the algorithms in WASP function. The results section details the analysis of these experiments, highlighting how aspects of the the polar writhe measure such as the local/non local and knot detection provide insight. These sections are aimed at providing a clear and easy to read guide to demonstrating how the WASP package can be used to provide insight into the evolving geometry of 3-D curves and ribbon structures such as those inherent in DNA structures. In addition, in the final section we show how WASP can be used to detect knot and belt-trick type deformations, again through an instructive example. Finally, we provide an supplement, which is of a much more technical nature and aimed at those readers with a specific interest in ribbon topology. It details how the original closed Călugăreanu theorem (the closed ribbon *Lk*-*Tw*-*Wr*) and the Fuller writhing quantities all naturally arise as part of the framework described in^16,26^. It is not necessary to read this supplement in order to use and interpret the WASP package, but it would be necessary in order to compare it to existing writhe calculation approaches.

## Introduction to the polar writhe

In typical applications, the *Lk* is a prescribed quantity (*i*.*e*. a fixed applied number of rotations of the structure in plectoneme experiments), and the twist *Tw* can be calculated as *Lk*-*Wr*. Thus, in this section, we focus on the definition of the writhing as the key quantity to be calculated. In^16,26^ the definition of writhing was given the name the *polar writhe* which we label *W*_*p*_ to distinguish from the pre-existing closed ribbon definition (the reason for this name will be clarified shortly). In what follows, we introduce the quantity *W*_*p*_ through a series of instructive examples.

### The Polar Writhe

The polar writhe^16^ is the sum of a local component (*W*_*pl*_) and a non local component (*W*_*pnl*_):

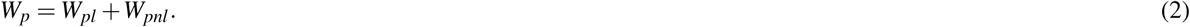

The twisted paraboloids shown in Fig. 2(a) provide an intuitive example of the behaviors tracked by these distinct components. The parabolas are described by the following formula:

**Figure 2.**
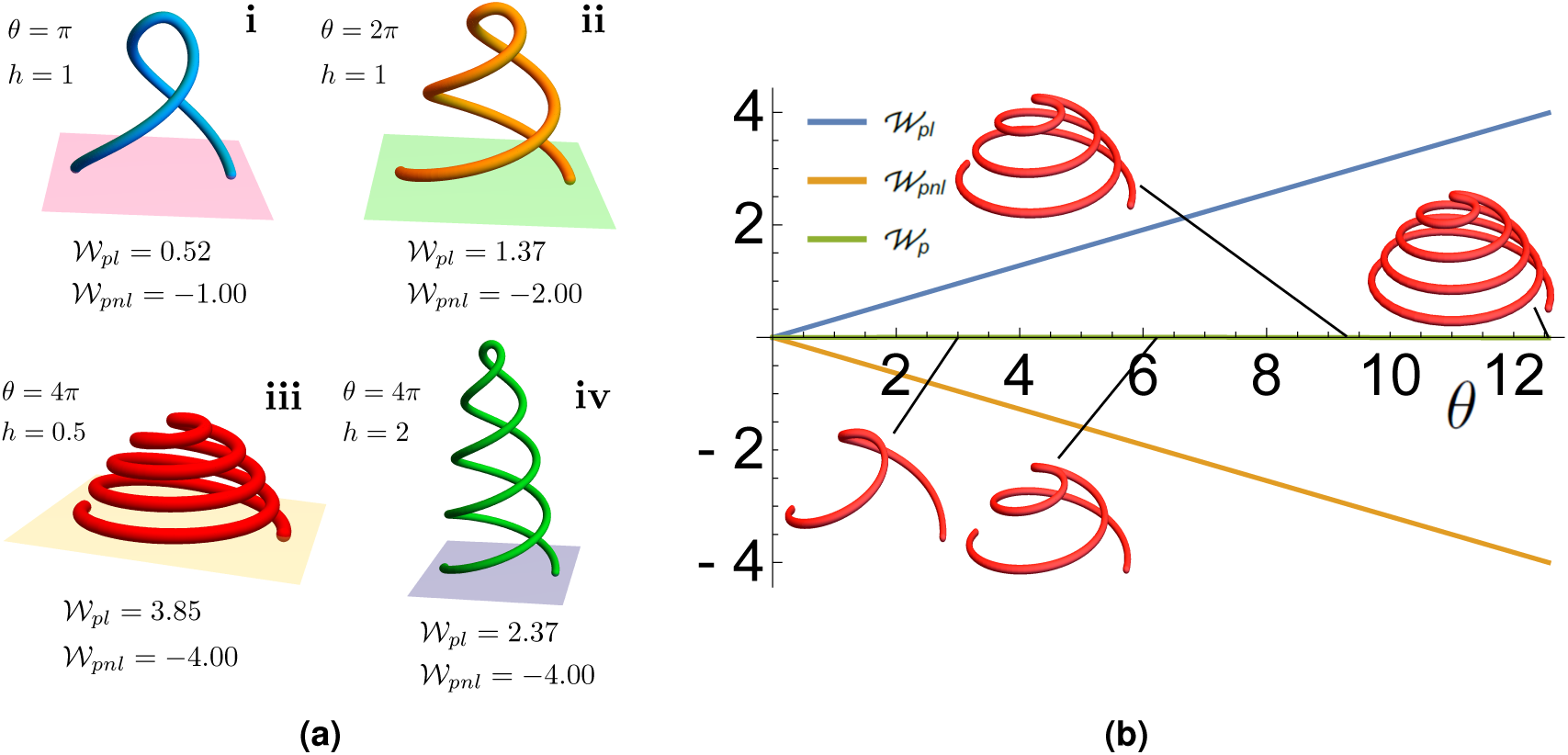
Twisted paraboloids use to highlight aspects of the polar writhe. Panel (a): paraboloids with varying heights *h* and winding angles *θ*. Panels (a)(i) and (a)(ii): paraboloids with equal height but different winding exhibit differences in both *W*_*pl*_ and *W*_*pnl*_ as a result of large-scale rotation and increased helical density. Panels (a)(iii) and (a)(iv): paraboloids with equal winding but varying height exhibit differences in *W*_*pl*_ only as the extent of buckling remains constant while helical density is varied. Panel (b): the polar writhe decomposition as a function of the winding angle *θ* for a parabola for which *W*_*p*_ = 0

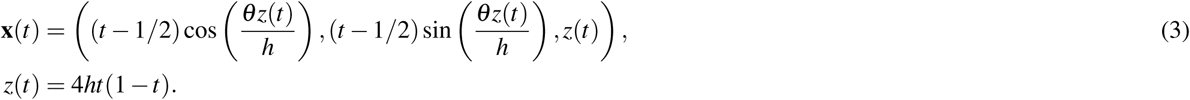

The parameter *θ*, the winding angle, determines the number of rotations applied to the parabola (*c*.*f*. Figures 2(a)(i) and (a)(ii)). The height parameter *h* determines the relative stretching of this coil (*c*.*f*. Figures 2(a)(iii) and (a)(iv)). **For this curve** *W*_*pnl*_ = *−θ/*2*π*. The factor of 2*π* indicates *W*_*pnl*_ measures the number of full rotations. The sign is due to the orientation of the curve as we shall explain shortly.

In the top row of Figure 2(a), *h* is fixed. In (a), *θ* = *π*, resulting in a single loop around the middle of the paraboloid. In this case *W*_*pnl*_ = *−*1. In (b), there is an additional looping and *W*_*pnl*_ = *−*2. The local component, which measures the local helical density of the paraboloid, is increased form (a)(i) to (a)(ii) due to additional winding at fixed height. In the bottom row we fix the winding (*W*_*pnl*_ = *−*4) but vary the curve’s height *h* to highlight the effect on *W*_*pl*_. As the height of paraboloid is increased from (a)(iii) to (a)(iv), the overall winding (and hence non local writhe) is unchanged, however, as the coil becomes increasingly less tightly wound, the local writhing decreases as its helical density is decreased. Note that in these cases the two components are generally of opposite sign, however, this is **not** a general property of the pair (*W*_*pl*_,*W*_*pnl*_). As shown in^16^ if one chooses *h ≈* 0.37 then *W*_*p*_ *≈* 0, irrespective of the choice of *θ* (see Figure 2(a)(ii)). We see the added value of the local/non-local decomposition providing the information about the increasingly tight helical coiling which is missing from the total sum.

#### Curve splitting

The idea behind the polar writhe calculation is that we split the curve into sections sharing a mutual vertical height. If we have a Cartesian coordinate system (*x, y, z*) where *z* is the height, then we split the curve at *turning points* for which the curve changes direction from upward to downward pointing: 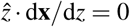. Examples of the parabolas (1 turning point, 2 sections) and a locally looped structure (2 turning points, 3 sections) are shown in Figures 3(a) and (b). In general, we label the subsections *i* of the curve that span heights 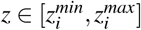 as **x**_*i*_. The local writhe quantifies the coiled geometry of each individual subsection and the non local writhe the mutual winding of all distinct pairs of sections.

**Figure 3.**
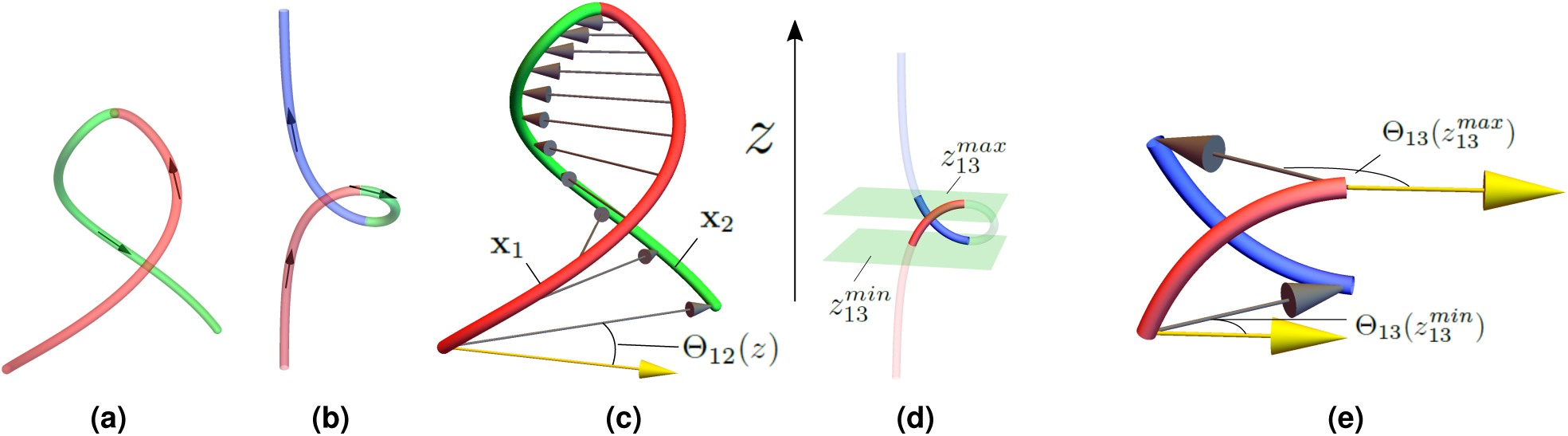
Illustrations of the non local writhe calculation. Panels (a) and (b) are curves split by their turning points. Their orientations are shown by arrows. Panel (a): the parabola’s turning point is at its peak and the two sections (red and green) which are partitioned by this turning point are shown. Panel (b): a looped curved which could represent the beginning of plectoneme formation. It has two turning points at the top and bottom of the loop. The curve is split into three sections, red, green and blue respectively, by these turning points. Panel (c): the two sections of the parabola **x**_1_ and **x**_2_ share a common mutual height *z ∈* [0, *h*]. At each height a vector is drawn from **x**_1_ to **x**_2_. As indicated, they make an angle Θ_12_(*z*) with respect to a fixed direction. The non-local polar writhe measures the rotation of this angle. Panel (d): subsections **x**_1_ and **x**_3_ of the curve shown in (b), which share a mutual *z* range between two planes 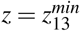 and 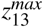. These subsections are shown in bold coloring. Panel (e): the *W* _*pnl*_ calculations for the mutual subsections shown in (d). The difference in the angles 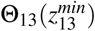 and 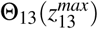 which characterises the non local writhing contribution form these two sections is depicted.

#### Local polar writhe

If the unit tangent vector of the curve **x** is **T** = d**x***/*d*s* (*s* the arclength of the curve), then:

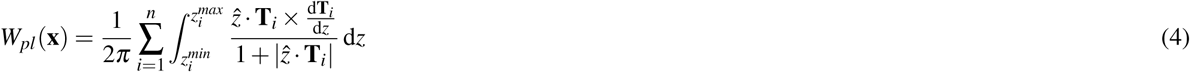

The numerator represents the rotation of the unit tangent vector around the vertical direction. It is positive if the curve coils in a right handed fashion and negative if left-handed. The denominator indicates (i): the rotation is given more weight if the curve is rotating relatively tightly (as with a small *h* parabola) and (ii): the modulus sign means that the orientation would be the same for a curve whether it points up or down, *i*.*e*. it only depends on the local helical chirality of the curve. The fact this rotation is measured around the 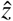 direction, which on a unit sphere of directions is the north pole, and that *W*_*pl*_ can be interpreted as an area on the unit sphere bound by the curve **T** and the pole, is the reason for the name the polar writhe (see^16^). However, this is not a crucial point in what follows and we do not mention it any further.

#### Non local polar writhe

Consider the parabola example. The curve is separated into sections **x**_1_ and **x**_2_ (as shown in Figure 3(c)). We define an angle Θ_12_(*z*) which is the angle made by the vector joining section **x**_1_ to **x**_2_ at a fixed height *z*, as shown in Figure 3(c). *W*_*pnl*_ measures the total number of rotations of this angle:

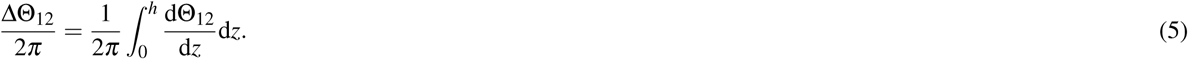

To ensure *W*_*p*_ forms part of an invariant sum with the twisting, the orientation of the curve sections is accounted for by multiplying by an indicator function *σ*_*i*_, with *σ*_*i*_ = +1 if the curve section **x**_*i*_ is moving upwards and *σ*_*i*_ = *−*1 if it is moving downwards. Also the calculation is counted twice, again this is necessary to form part of the invariant sum *W*_*p*_ + *Tw*. Thus for the parabola

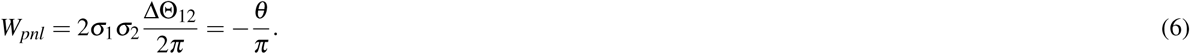

For the looped curve shown in Figure 3(b) we follow a similar procedure for the pairs (**x**_1_, **x**_2_), (**x**_1_, **x**_3_) and (**x**_2_, **x**_3_). The only extra complication is that the curve sections do no share the same mutual *z* ranges and the winding is only measured for subsections of the curve which share the same mutual *z* range 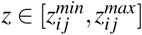, as indicated in Figure 3(d). The non local writhe for the pair (**x**_1_, **x**_3_) would be:

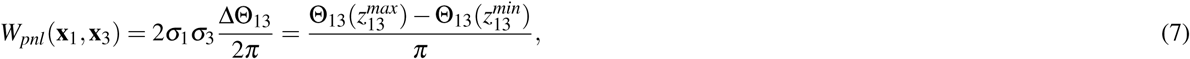

as indicated in Figure 3(e). Note the product of *σ*_1_*σ*_3_ here is positive as both sections have the same vertical orientation. The total *W*_*pnl*_ of this curve would be:

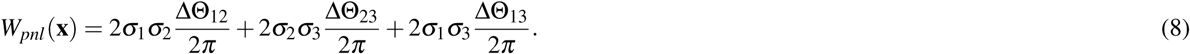

In general *W*_*pnl*_ is just the mutual winding of all subsections of the curve which share a mutual height range. In the general case where the curve **x**_*i*_ has *n* sections (and *n−* 1 turning points) the non local writhing is calculated as

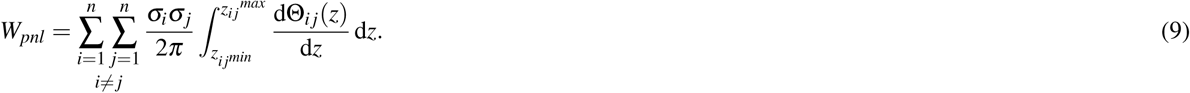

#### Looped local to non-local transition

We consider a second illustrative example: a curve transitioning from a locally helically coiled curve to a looped curve, the first part of plectoneme formation. The looped curve being the curve shown in Figure 3(b) used as an example above. This transition is described by a set of curves which are equilibria of elastic rod models:

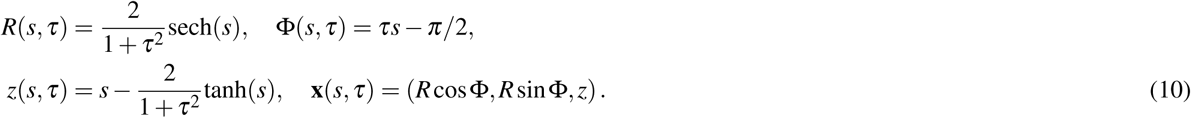

Here *s ∈* [*−*5, 5] is the curve’s arclength and *τ* a parameter which acts to twist the tube as it is decreased, here from 3 to 0.2. As the tube is twisted its axis transitions from a curve with a helical perversion around its centre, see Figure 4(i), to a looped curve indicative of the beginning of plectoneme formation as shown in Figure 4(ii). One can obtain this behaviour by steadily twisting a cable under tension; that they are are equilibria of an elastic tube model is shown in^38^, and thus they represent the deterministic counterpart of a worm-like ribbon model *e*.*g*.^7,8^. In Figure 4 we see its *W*_*p*_ values as *τ* is decreased from 3 to 0.2 in 200 evenly spaced steps. The polar writhe *W*_*p*_ is seen to steadily increase as the curve becomes at first increasingly helically coiled at its centre and then increasingly looped. For curves 1-138 all the contribution is local due to the helical kinking of the curve around its centre (*W*_*pnl*_=0). After this the loop begins to form and the curve develops non-local writhing (*W*_*pnl*_ *>* 0). What is interesting is the dramatic inter-conversion of local and non-local writhing which occurs whilst the sum changes smoothly through this transition, indicating that the heilical coiling becomes distorted to form the loop. Again we see the local/non local decomposition providing significant extra information which characterizes the geometrical development of the curve.

**Figure 4.**
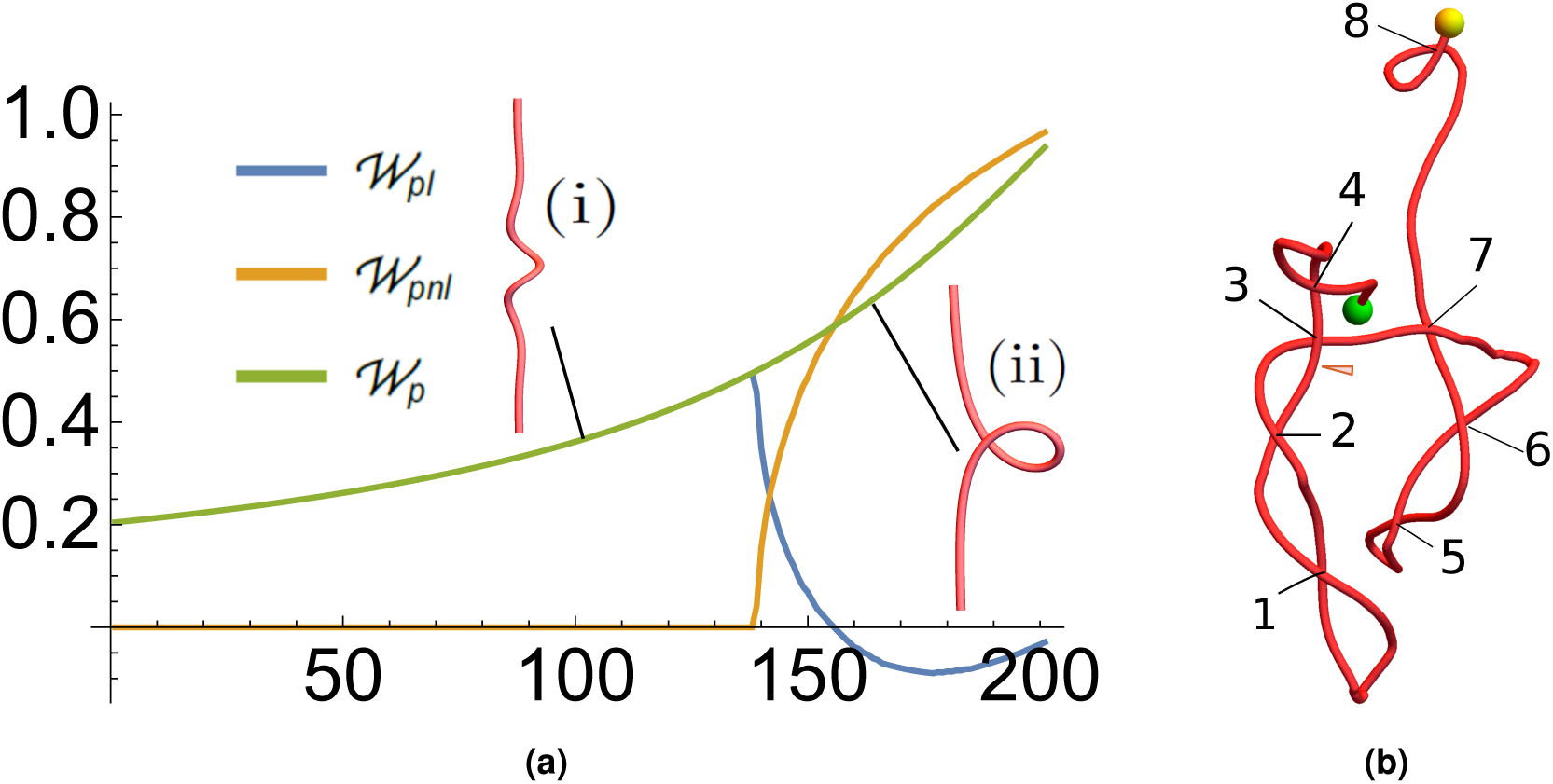
Polar writhe calculations of plectoneme geometries. Panel (a) *W*_*p*_ values of a loop forming curve deformation. Panel (b), curve with significant plectoneme structure. The individual loops present are marked.

#### Non local writhing detecting plectoneme formation

An example of a highly plectonemed curve is shown Figure 4. Marked on the curve are 8 clear crossings indicating looped geometry. Since we have seen in the parabola example (Figure 2) loops typically have a *W*_*pnl*_ value of *±*1 we should expect the non local writhing of this curve to have a magnitude of around 8. In fact for this curve *W*_*p*_ = 8.88, *W*_*pnl*_ = 8.55, *W*_*pl*_ = 0.33 (all to 3.s.f). So we can infer that for highly plectonemed curves *W*_*p*_ will be largely dominated by non-local writhing and will roughly count the number of loops of the plectoneme.

Finally, for the interested reader an additional set of example calculations, including local-non-local decompositions of example curves can be found in Chapter 4 of^39^ and^16^, and examples of elastic rod equilibria in Chapter 6 of^26^. We will shortly turn our attention to calculating this quantity for the output of numerical supercoiling experiments.

### Self crossing detection

Both the linking *Lk* (the net winding, given by the sum *W*_*p*_ + *T*_*w*_) and the polar writhing *W*_*p*_ change by a a value of *±*2 if the ribbon intersects itself (the twist changes continually). This would for example be relevant in DNA models for topoisomerase action. In^26,36^ this fact was used to detect whether solution branches for an elastic tube model were physically valid. A series of evolving elastic equilibria were obtained using a continuation method, when the values of *W*_*p*_ jumped by *±*2 the could be automatically ruled out as not physically accessible (as elastic tubes cannot self intersect). Of course in DNA modelling we typically expect this to be impossible due to localized repulsion of the polynucleotide chains.

## Methods

### Numerically simulating DNA tweezer experiments

Plectonemic DNA structures were generated via coarse-grained molecular dynamics simulations of magnetic/optical tweezer experiments. 600 base-pair linear DNA helices were constructed with three different net-windings: *Lk* = 40 (*≈* 24*°*per turn) (underlinked), *Lk* = 60 (*≈*36*°*per turn) torsionally relaxed, and *Lk* = 80 (*≈*48*°*per turn) over-linked. Their evolution was simulated using the oxDNA2 model^40^. To simulate tweezers experiments, each helix was first aligned such that its axis was along the 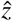 direction. Anchor restraints were placed on the bottom five base-pairs of each helix to permanently fix their position, and harmonic restraints were applied to the top five base-pairs of each helix to allow the respective base pairs to move only in the 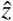 direction and also prevent the DNA from shedding its winding by simply untwisting. In addition to these restraints, a range of extension forces (0pN, 2pN, 4pN, 6pN, 8pN) were applied to each helix in separate simulation by pulling the top five basepairs in the +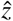 direction. Each helix was simulated for a total production run of 1.2×10^8^ time steps using the oxDNA2 model with a salt concentration of 0.5 M. We only detail three specific cases below for brevity. These cases are chosen as they highlight the critical characteristics of the polar writhe measure as applied to these numerical supercoiling experiments

### Atomistic molecular dynamics simulations of supercoiled DNA minicircles

108 base-pair minicircles were constructed with *Lk* = 14 (Δ*Lk* = +4) to promote conformational variety/structural buckling predicted by previous experimental/simulation explorations of minicircle dynamics^41,42^. Two base-pair sequences, (AA)_27_(AT)_27_(AA)_27_(AT)_27_ and (GG)_27_(GC)_27_(GG)_27_(GC)_27_, were explored as well. Simulations were performed in explicit solvent using AMBER with a modified DNA BSC1 force field to correct for the circularized DNA and Smith and Dang ion modifications for Na and Cl ions^43–45^. DNA minicircles were solvated in a TIP3P rectangular water box with a 15.0Å buffer. Following solvation, the appropriate cosolute (spermine) was placed along with the DNA within the water box and the entire system was neutralized with Na+ ions. Initial minimization was performed on the cosolute and water molecules with a 10 kcal/mol restraint placed on the DNA residues. After the initial minimization, restraints were removed from the DNA and the entire system was allowed to minimize. Following the minimization phase, restraints were reapplied to the DNA residues and the system was heated from 100K to 300K. Consequent to heating, restraints were progressively removed from the DNA residues over a period of *≈*4ns. MD was performed without restraints on the equilibrated system for 100ns for each trial in the NVT ensemble. The number of spermine molecules in each simulation varied between trials to explore different ranges of cosolute concentration.

### Obtaining a DNA axis

To calculate *W*_*p*_ it is first necessary to generate an axis for the DNA molecule. In order to properly characterize the geometry of a given DNA structure, it is important to generate this axis curve in a manner such its total curvature is minimized. Failure to do so may result in the misrepresentation of the DNA geometry due to excess local writhe introduced to the axis curve as a result of the helical nature of the DNA backbone strands. Simplistic axis curve formulations such as the axis curve obtained by taking the midpoint of the C1’ atoms located on opposite sides of each base-pair along the helix result in coiled axes that exhibit the same helical periodicity as their encompassing backbone strands. To avoid such misrepresentation, the WASP package uses an implementation of the WrLINE axis curve method developed by Sutthibutpong, Harris and Noy^46^ which effectively smooths the contour of the axis curve, eliminating the unwanted “coiling” and helical periodicity exhibited by alternative methods that generate unwanted excess local writhe. Full details regarding the WrLINE formulation can be found in^46^, but key features of the formulation are summarized below maintaining similar notation as in^46^.

Unlike a basic midpoint method in which an axis is obtained by simply calculating the midpoint *m*_*i*_ of the pair of C1’ atoms (*r*_C1’,A_ & *r*_C1’,B_ - see Fig. 5) on opposite sides of a corresponding basepair *j*:

**Figure 5.**
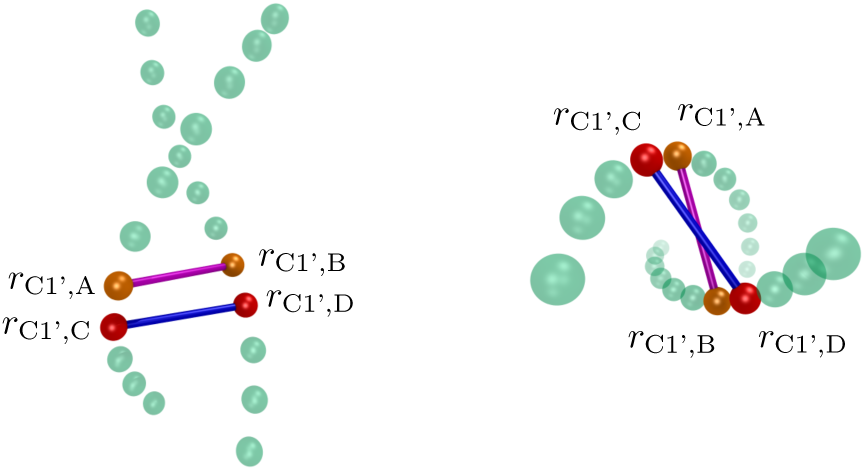
Left: Side perspective of backbone atoms along a helical fragment of DNA. Beads **r** represent C1’ atoms along the DNA backbone. Right: Top view of backbone atoms along the helical fragment.

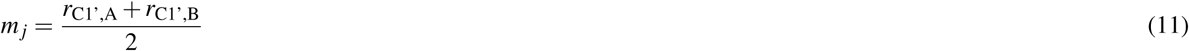

for each base-pair along the DNA helix, the midpoint *r*_*i*_ of each “dinucleotide” (2 base-pair) step along the DNA helix consisting of the two pairs of C1’ atoms on opposing sides of a set of two consecutive base-pairs *j* & *j* + 1 on the helix is calculated as:

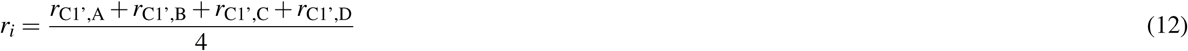

The remainder of the WrLINE method can then be roughly summarized by the following procedure:

1. Given a dinucleotide midpoint *r*_*i*_, calculate the base-pair step twist *θ*_*i*_ from the dinucleotide step corresponding to *r*_*i*_
2. Determine the number of base-pairs (2*m*) required to complete a full helical turn around the base-pair step corresponding to *r*_*i*_ (i.e. *m* base-pairs above *r*_*i*_ and *m* base-pairs below *r*_*i*_.)
3. Calculate a weighting factor *w* related to the sum of the base-pair twists *θ*_*i−*1_, *θ*_*i*+1_, *θ*_*i−*2_, *θ*_*i*+2_ … throughout the helical turn around *r*_*i*_.
4. Use *r*_*i*_, *m*, and *w* to determine an point on the axis curve *h*_*i*_.

The above procedure is then repeated for each midpoint *r*_*i*_ to obtain a full set of axis points *h*_*i*_ that represent the DNA helical axis. Note that the summary of the WrLINE method provided above has been included solely for the purpose of familiarizing the reader with the axis curve method utilized in WASP as well as to highlight the intricacies of determining a proper helical axis. The reader is strongly encouraged to refer to^46^ for a comprehensive treatment of the WrLINE method.

It is important to note that the WrLINE method is intended to be used primarily with closed DNA structures such as DNA minicircles. As such, the algorithm requires that there exist a full helical turn of DNA around each point *r*_*i*_ as specified above. This requirement is problematic in the case of linear DNA helices where a full helical turn of DNA cannot exist around points *r*_*i*_ near the ends of the helix. In order to circumvent this issue, two options are available:

1. Treat the DNA structure as though it were closed (joined at the ends) despite the fact that it is, in reality, not. This allows the WrLINE method to be utilized verbatim as outlined in^46^, but will result in a number of undesirable stray axis points generated in arbitrary positions external to the DNA structure. The number of stray axis points will be proportional to the extent of supercoiling of the helix (how many base-pairs are required to make a full helical turn) and can be safely, manually deleted via knowledge of the system/visual inspection. These stray axis points will always constitute the “ends” of the unaltered axis as they are a product of end effects produced by the WrLINE algorithm. This option is the most general, but should be utilized with extreme caution.
2. Start the WrLINE method on the first points *r*_*i*_ such that one full helical turn of DNA can be formed around the respective points *r*_*i*_ on each end of the DNA helix (satisfying the criteria for the calculating the Θ_*m*_ parameter^46^) as determined prior to beginning the axis curve computation.

At the time of writing, options 1 and 2 are implemented in WASP via the **deleteatoms** and **autodelete** arguments respectively. In either case, the total number of resulting axis points will be less than the total number of base-pairs when evaluating open DNA structures using the WrLINE method properly.

### Calculating *W*_*p*_

The WrLINE algorithm yields a discrete curve 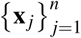. The algorithm for calculating *W*_*p*_ for this curve is as follows:

1. Split the curve into *n* sections **x**_*i*_ by locating its turning points. This is done from the set of tangent vectors **T** _*j*_ = **x** _*j*+1_ − **x** _*j*_ and use of tricubic interpolation.
2. For each section **x**_*i*_ calculate *W*_*pl*_(**x**_*i*_) using equation (4). The WASP package uses a modified version of Simpson’s rule^47^.
3. Identify the range of mutual overlap of all pairs of sections (**x**_*i*_, **x**_*k*_), the WASP package uses a simple binary search algorithm.
4. Use equation (9) to calculate *W*_*pnl*_ (a branch cut tracking method is used to track full windings).

This is implemented in C++ for the WASP package, but the main routines can be accessed by a python interface.

In addition, for comparison to existing calculations, we calculate the writhe as defined by:

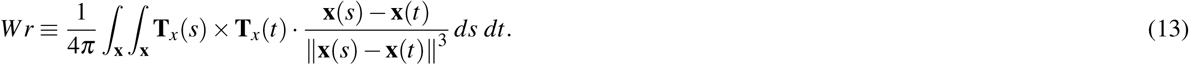

using the algorithm specified in^48^. This is the expression for writhing used in the closed ribbon Călugăreanu theorem (applied here to open curves). As discussed in the introduction, this measure has been used in open ribbon studies to characterise the writhing geometry of the ribbon’s axis and we present the results here for comparison. We do not present any results using closures. It was demonstrated in both^16^ and^26^ that there is always a closure for which the closed ribbon structure will have the exact same writhe value as *W*_*p*_ (in fact it is the typical stadium closure), a numerical demonstration of this fact is also detailed in^26^. Further it was shown that the closure can account for a significant proportion of the calculation, hence obscuring the interpretation. This last fact is the reason the polar writhe local/non-local decomposition is a preferable measure for quantifying the ribbon’s evolving geometry.

## Results

### Undertwisted Helices (*Lk* = 40)

Undertwisted helices exhibited relatively modest writhing dynamics throughout their trajectories. In contrast with torsionally relaxed helices, helical strain due to under-twisting results in the formation of small kink structures in the curve as indicated in Figure 6(a). We detail the results here of two particularly pertinent cases here, a weak 2pN stretching force and a stronger 8pN force.

**Figure 6.**
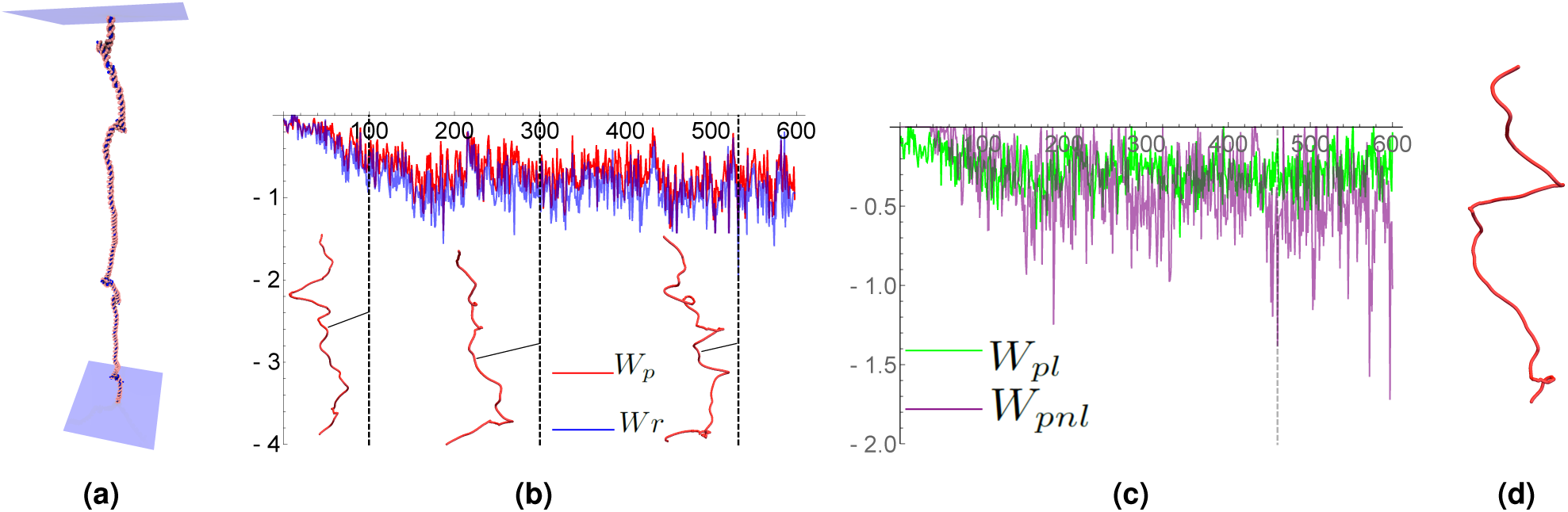
Results from simulations of an *Lk* = 40 DNA helix (undertwisted) placed under a 2pN extension force. Small kinked structures occasionally form along the length of the structure, as shown in panel (a). Shown in panel (b) are writhe time series plots. The *W*_*p*_ values are shown in red and the *Wr* values in blue. The *Wr* values are consistently larger in magnitude but the variations in magnitude follow a very similar pattern. Example axis curves from the time series are shown, they correspond to the times marked by the dashed vertical lines. The curves width/height ratios are 0.3, chosen for clarity (0.1 would be the actual ratio). Panel (b): a comparison of *W*_*pl*_ and *W*_*pnl*_ for the *W*_*p*_ calculations shown in panel (a), there is a mixture of local *W*_*pl*_ and non local writhing values *W*_*pnl*_ of roughly equal value. Some spikes in *W*_*pnl*_ were found to arise from temporary loop formation, the dashed line indicates one such example whose curve is shown in panel (d). The spike in *W*_*pnl*_ can be seen to correspond to the tight loop towards the bottom end of the curve.

The 2pN results are shown in Figure 6(b). The values of both writhe measures *Wr* and *W*_*p*_ settle at values at about *−*1, a twentieth of the applied (under) rotation. The *W*_*p*_ value is consistently lower than the *Wr* value, although not by a substantial amount. As indicated in the figures there is no plectoneme formation, however, often partially looped sub-sections form. The local/non-local decomposition of *W*_*p*_ represents this mixture of helical (local) distortion and non-local coiling Figure 6(c). There are occasional spikes in the non-local writhing (relative to the local value). One such example is shown to arise from a tight loop formation as shown in Figure 6(d).

The 8pN results are shown in Figure 7(a). The additional force further restricts the writhing of the curve and the magnitude of both quantities is typically bound between [*−*0.5, 0.5]. The value of *Wr* is typically significantly larger than its *W*_*p*_ counterpart. We verified that the ratio |*W*_*p*_ *− W*_*r*_| */*(|*W*_*p*_| + |*W*_*r*_ |) was larger than 0.5 for over a third of the calculations (*>* 50% difference). In Figure 7(b) we see that for vast the majority of the curve’s geometries there is only local writhing, a clear difference from the 2pN case indicating the force is acting to restrict loop formation. The non-local writhe values arose when small kinks in the structure developed, although again these are significantly restricted by comparison to the 2pN case.

**Figure 7.**
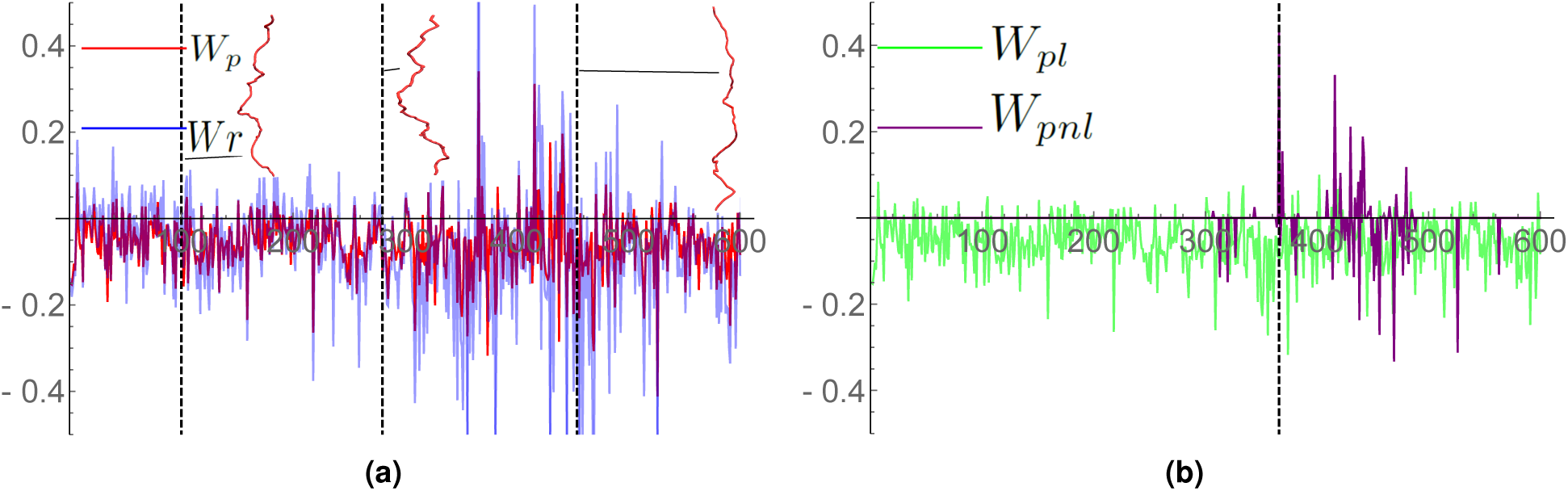
Writhe time series plots for an undertwisted DNA molecule (*Lk* = 40), subjected to an 8pN pulling force. Panel (a), a comparison of *W*_*p*_ and *Wr* for the time series. The *W*_*p*_ values are shown in red and the *Wr* values in blue. The *Wr* values are consistently larger in magnitude. Example axis curves from the time series are shown, they correspond to the times marked by the dashed vertical lines. The curves width/height ratios are 0.3, chosen for clarity (0.1 would be the actual ratio). Panel (b): a comparison of *W*_*pl*_ and *W*_*pnl*_ for the *W*_*p*_ calculations shown in panel (a). For the significant majority of curves there is only local writhing.

### Overtwisted Helices (*Lk* = 80)

Overtwisted helices exhibit dramatic writhing as a result of high torsional strain on the DNA helix. As a result, helices contracted over time in the 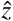 direction by forming a pair of plectonmeme structures (Figure 8(a)). Time series plots of *W*_*p*_ and *W*_*r*_ are shown in Figure 8(b) for a helix experiencing no extension force. There is a steady increase in writhing up to nearly 10 in both cases as the curve forms the plectoneme structures, this is half the applied over-rotation so there has been a significant conversion of (over)twisting into writhing. We note the two measures *W*_*p*_ and *Wr* are nearly identical unlike in the under-twisted case. The reason for this is discussed in more detail in the supplement, it suffices to state here that both assign the same value to the loops in the plectoneme structures and these dominate the axis curve’s geometry.

**Figure 8.**
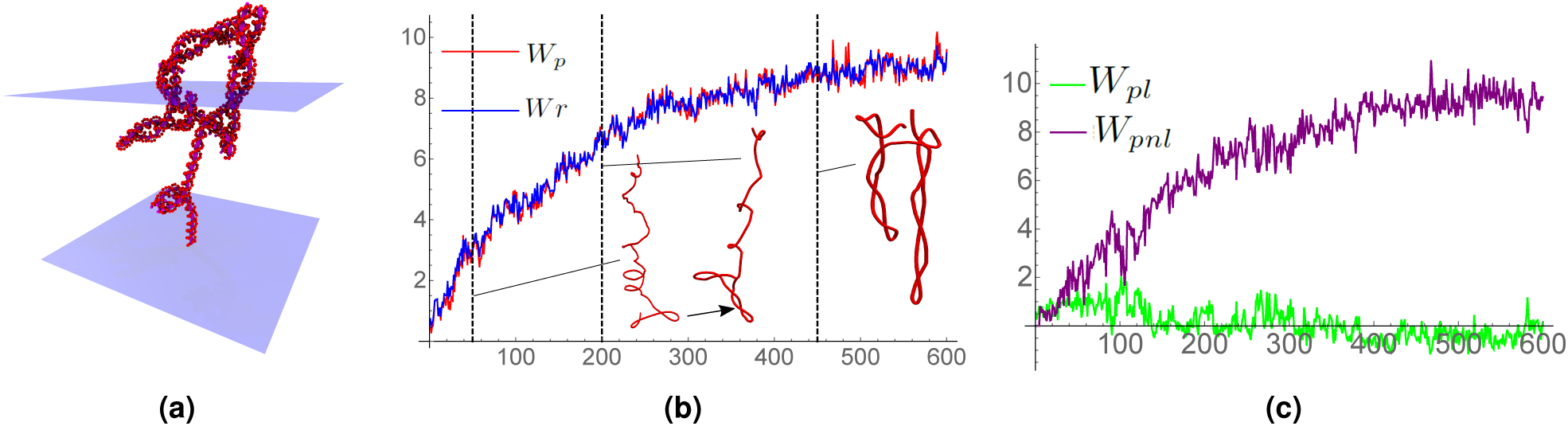
Results from simulations of a *Lk* = 80 DNA helix (over twisted) experiencing no extension force. In panel (a) we see the formation of significant supercoiling. In panel (b) we see the development of this supercoiling is reflected by the steady increase in both writhing measures. Example axis curves from the time series are shown, they correspond to the times marked by the dashed vertical lines. The curves width/height ratios are respectively 0.5, 0.4, 0.3 and chosen for clarity (0.1 would be the actual ratio). Panel (c) shows the local/non-local decomposition of *W*_*p*_. Except in the initial stage the non local writhing is dominant and it increases over time as the curve forms increasing numbers of plectoneme type loops.

The local/non-local decomposition is shown in Figure 8(c). Generally, except in the initial stage of simulation, there is far more non-local than local writhing, again due to the plectoneme formation. The first curve shown in Figure 8(b) at time step 50 has significant helical looping in its upper part, nearly the lower end of the curve, however, we see the initial formation of the first plectoneme, this contributes non-local writhing. As indicated in 8(c) there is a reasonable balance of local and non-local writhing reflecting this two part geometry. The looped section has formed a plectoneme by time step 200, as shown in Figure 8(b). By this stage the non local writhing is dominant 8(c). The final curve shown in Figure 8(b) shows two plectonemes, the second developed from one of the small loops in the upper half of the curve at step 200.

### DNA Minicircles

To further highlight the utility of *W*_*p*_, we have included sample analyses of two DNA minicircles (Figure 9). Both trajectories represent 108 base pair *Lk* = 14 minicircles exposed to a 10 mM spermine cosolute. The minicircle in 9(a) has the base pair sequence (GG)_27_(GC)_27_(GG)_27_(GC)_27_ and the minicircle in 9(b) has the sequence (AA)_27_(AT)_27_(AA)_27_(AT)_27_. In both trajectories, *W*_*pnl*_ is observed to gradually increase throughout the beginning of the trajectory and then fluctuate around an average as the minicircle system reached an equilibrium state. In these examples, the measurement of *W*_*pnl*_ reflects the gradual formation of loops/buckling points within the minicircle structures caused by spermine bridging. Spermine molecules adsorb to the backbone of the DNA minicircles early in the trajectories and gradually bridge together adjacent tracts of the DNA minicircles over time resulting in the formation of loops/buckling points within the minicircle structures as shown in Figure 9(c). The observed bridging dynamics are in accordance with results from previous studies as sequence dependent interactions have been observed between spermine molecules and AT/AA rich groups of linear DNA helices in which spermine molecules were observed to adsorb parallel to the DNA backbone and bridge together adjacent DNA helices^49^.

**Figure 9.**
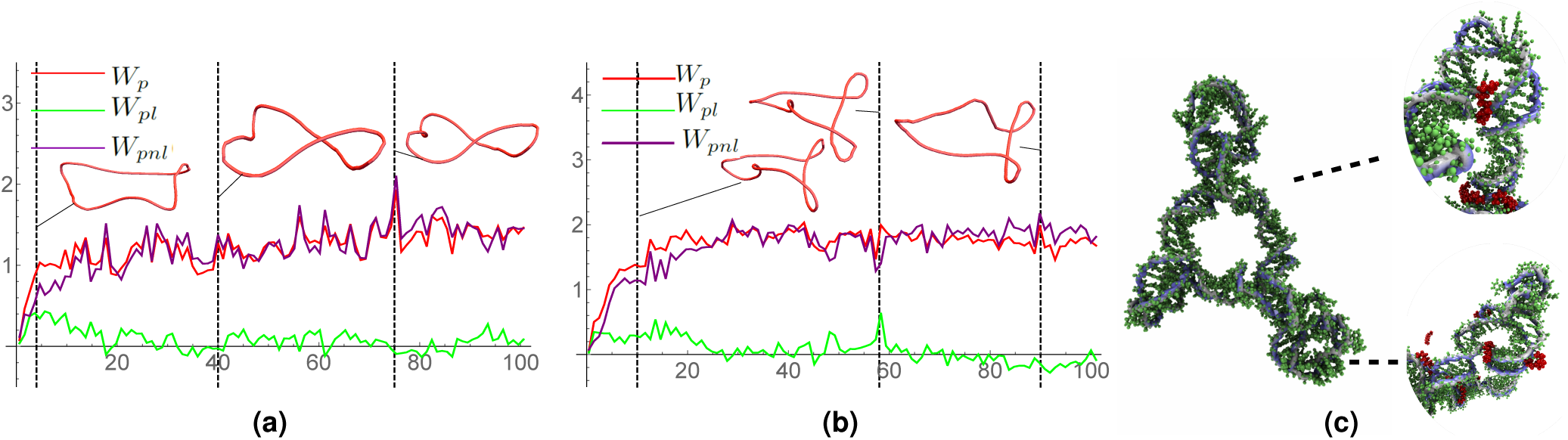
Polar writhe calculations for two DNA minicirlce cases. Panel (a) indicates the writhing evolution of a 108 base pair *Lk* = 14 minicircle with a (GG)_27_(GC)_27_(GG)_27_(GC)_27_ sequence exposed to a 10mM spermine cosolute concentration. Panel (b) indicates the writhing evolution of a 108 base pair *Lk* = 14 minicircle with an (AA)_27_(AT)_27_(AA)_27_(AT)_27_ sequence exposed to a 10mM spermine cosolute concentration. Panel (c) depicts spermine bridging in an (AA)_27_(AT)_27_(AA)_27_(AT)_27_ minicircle. Spermine molecules are shown in red.

As shown in 9(a), the axis of the GC/GG minicircle forms a figure 8 structure with a smaller loop occasionally forming at a consistent location on the curve. The *W*_*p*_ value is generally dominated by the non-local component, with variations being linked to the development (and reduction) of the smaller localised loop. In panel (b) it is shown that the AT/AA minicircle quickly forms two clear loops giving a *W*_*p*_ value which oscillates around 2. Again there is small localised loop which variable develops/recedes. At one point *t* = 49 marked with a dashed line we see a relatively large spike in the local writhing corresponding to the loop localising and forming a tight local helical loop.

For both the AT/AA and GC/GG minicircles, the small localised loops that develop/recede within the minicircle structures are caused by spermine molecules that “jump” around the backbone of the DNA in various locations. These spermine molecules intermittently adsorb to the DNA backbone, bridging sections of the minicircle together which ultimately results in the formation of small loops that recede once the spermine molecules detach from the backbone and move to a new location. It is interesting to note that the non-local component of *W*_*p*_ serves as a direct indicator of this phenomena, fluctuating directly in response to the aforementioned spermine/loop formation dynamics.

## Writhe star and winding classes

A complication with open ribbons such as extended DNA structures is the possibility of (un)knotting and belt-trick type deformations, such as those shown in Figure 10^17^. These are classes of shape changes in which a section of the ribbon structure loops over one of its end points (as in Figure 10(b)-(d)). In such cases both *Lk* and *W*_*p*_ can change by values of *±*2^26^. Thus, when one prescribes a fixed value of *Lk* it is only constrained up to an integer, but critically that integer change can only occur when an “over-the-top” deformation occurs. For many applications this will not be possible, for example DNA structures in restricted environments and magnetic bead experiments where the bead itself is barrier. However, other potential applications of the writhing such as a measure of the development of tertiary structure in protein folding may need to consider such deformations. Prior and Neukirch^26^ considered how over-the-top deformations affect the polar writhe formulation, we briefly introduce that framework here through some examples of of how this integer change can be tracked. This facility is built into the WASP package. **We should stress that if this over-the-top transition is prevented then one simply needs to use the** *W*_*p*_ **calculations as described in the previous sections and what follows is unnecessary**.

**Figure 10.**
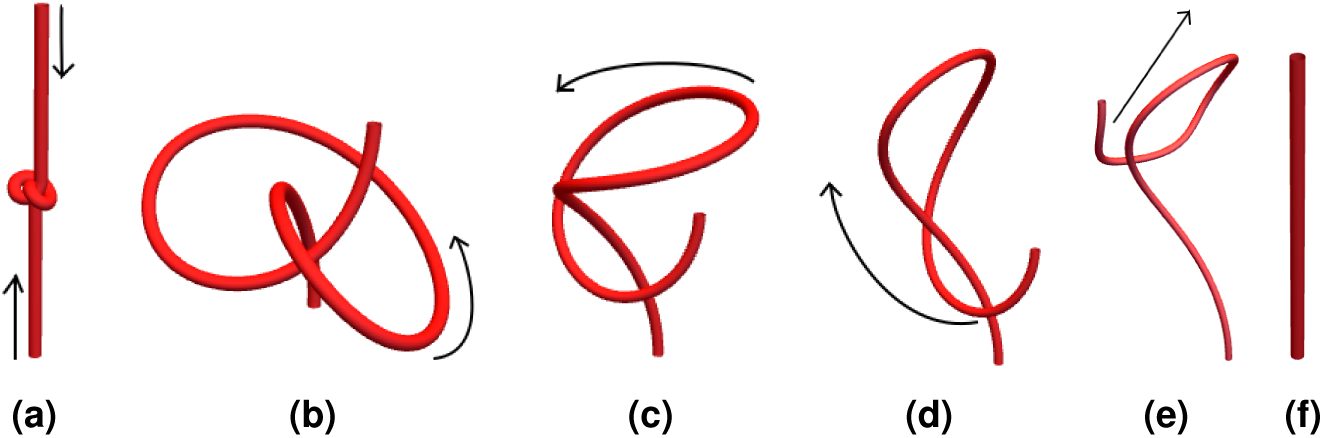
An “over-the-top” unknotting deformation. An initially tight knot shown (a) is relaxed (a)*→*(b). A section of this knot is then allowed to loop over the top of one end of the curve, (b)*→*(d). The curve is then pulled straight to an unknotted straight line (d)*→*(f).

### End angles and pulled-tight topology

We can think of the two ends of an open ended ribbon as being bound between planes. If any section of curve pases through the end of these planes it makes an angle with the end point as shown in Figure 11(a). This angle forms part of the non-local writhing calculation equation (9). As the curve loops over the top of the end point this angle increases Figure 11(a)-(d). This change would affect both *W*_*p*_ and *Lk*. If the curve then comes back down on the other side of the end point this angle will have gone through a change of 2*π* and since this is counted twice and change in *±*2 to both *W*_*p*_ and *L*_*k*_. So overall the change of *±*2 is detected.

**Figure 11.**
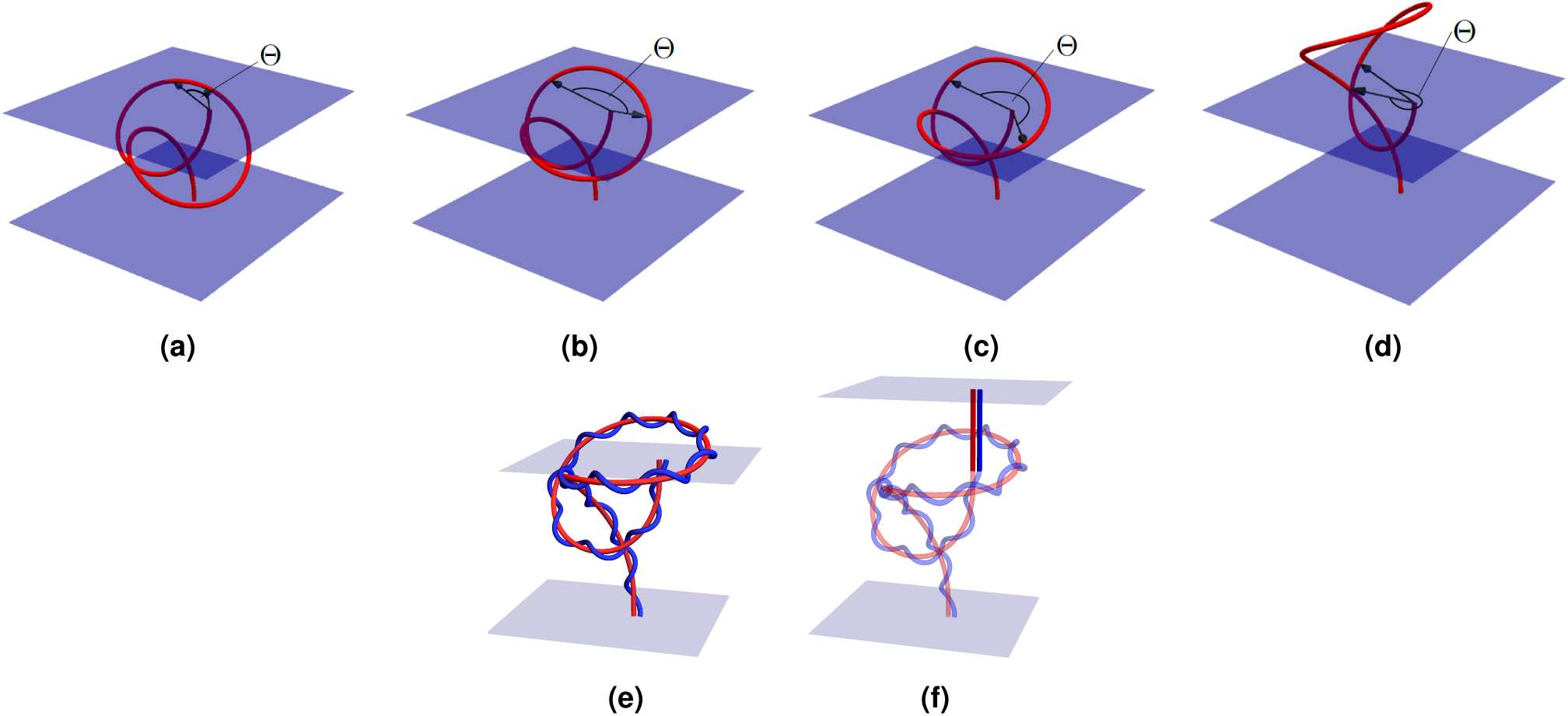
Illustrations of the loss in writhing which occurs to a section of the curves interior looping over one of its end points and the extension used to make this loss an discontinuous jump which tracks the pulled tight topology of the ribbon. Panel (a), an interior section of the curve rises above one of the planes containing the curve’s endpoints. The angle Θ represents the difference of two contributions to the *W*_*pnl*_ calculation which involves angles made with the curves end point. In panels (b) and (c) the curve section rises higher and the angle Θ increases. In panel (d) the angle is nearly 2*π* and the curve section has (just) passed directly over the top of the curve’s end. Panel (e) is a ribbon corresponding to one of the curves in Figure 11, a section of the curve is in the plane above the ribbon’s end points. Panel (f): the planar extension to the ribbon is shown. Now the whole ribbon structure is bounded between two planes. The extension is composed of planar curves with no twisting.

However, this means both *W*_*p*_ and *Lk* would be changing continuously (although still such that *Lk − W*_*p*_ = *Tw*). **This change does not happen if the curve is bound between the two end points where** *Lk* **is a fixed quantity**. For magnetic field applications where the end planes represent the edge of the system, *i*.*e*. the sun’s surface^50,51^ or the edge of the measurement domain in a plasma experiments^52^, evaluating this angle is crucial. However, for plectoneme formation and protein topology type applications it is important that the section of curve is not ignored from the calculation. Prior and Neukirch^26^ derived a variant of the *Lk* = *W*_*p*_ + *Tw* formula, with quantities 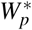 and *Lk*^*\**^ which in effect ignore these end angles and just register a jump in both *W*_*p*_ and *Lk*. More precisely it was shown one could always extend the ribbon with straight sections (with no *Tw*) such that the ribbon remains bound between two planes. This adds no local writhing or winding so in effect remains a measure of the topology of the original unextended ribbon, except in one crucial aspect, when the curve/ribbon passes over its end end point. Then the ribbon self intersects and there is a jump in *±*2 in both *Lk* and *W*_*p*_ and thus *Lk* is fixed up to integer multiples of 2. This has the following intuitive interpretation. If we imagine taking ribbon’s axis curve end points and pulling them directly apart, then a section of curve passing over the end point marks the point in its deformation where the curve would change its “pulled tight state”, *i*.*e*. Figure 10(a) to (f). For those readers familiar with the literature of artificial closures there would be the same jump when the curve intersects its closure.

In summary, the sum

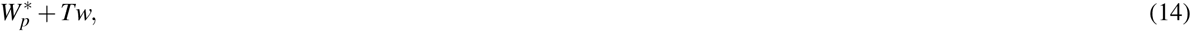

is invariant under all deformations which forbid curves passing over their end points, and changes by values of *±*2 when these over-the top deformations occur. This integer change can be used to indicate the ribbon’s pulled-tight topology (the shape obtained by pulling its end directly apart) has changed.

### Examples

First we note that the parabola and looped example calculations shown in Figure 2 and Figure 4 would be identical. As would the undertwisted DNA results shown in Figures 6 and 7.

#### Knot undoing

The curves depicting the undoing of a knot in Figure 10 are part of a continuous set of deformations. This deformation was discretized into 360 steps. In the first 100 steps, Figure 10(a) to Figure 10(b), are simple in that the shape of the knotted (trefoil) curve does not change but its size with respect the the straight end extensions is increases; in short the shape of the curve is effectively unchanged. From curve 101 to 260 the knot is undone as a section of the curve loops over its top end point, Figure 10(b) to Figure 10(d). Then from step 261 *−* 360 the curve is pulled straight 10(d) to Figure 10(f). The net effect is the undoing of a knot.

The values of *W*_*p*_, 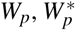 and *Wr* (given by (13)) are shown in Figure 12. For the first 100 steps both *W*_*p*_ and 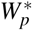 are identical and unchanged, correctly representing the fact the knot’s shape is essentially unchanged and the corresponding net winding would not change (assuming as always the ends of the ribbon are preventing form untwisting). The *Wr*, calculated via the Gaussian integral, equation (13), however, does change. The reasons for this change somewhat technical and are discussed in the supplement, it suffices to note here that it is not characterizing the knots unchanging shape and the sum *W*_*r*_ + *Tw* would **not** be a fixed quantity.

**Figure 12.**
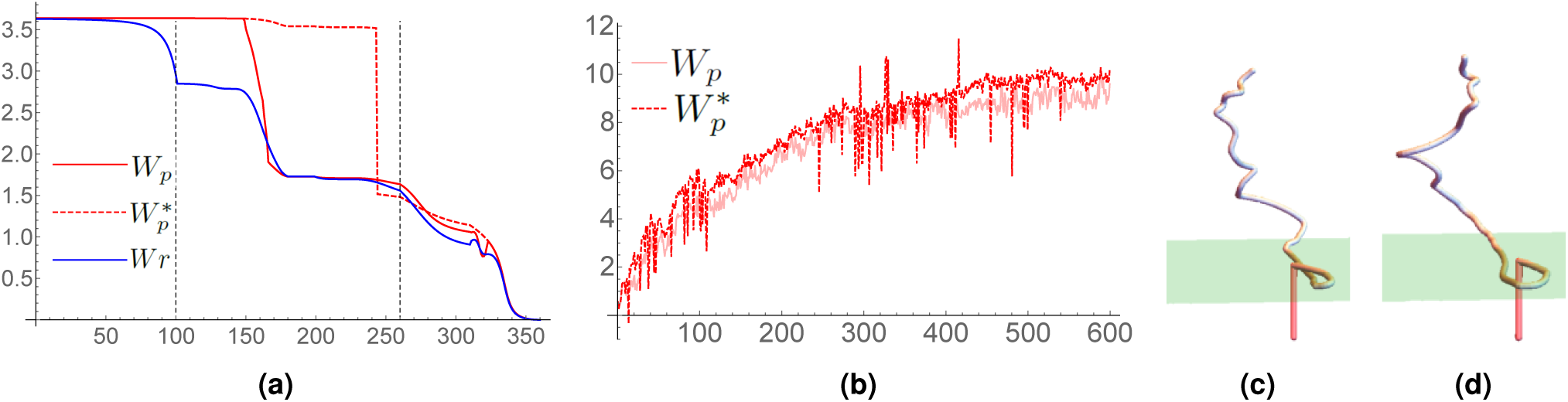
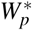 calculations. Panel shows (a) writhing values of the curves shown in Figure 10. Panel (b), a comparison of *W*_*p*_ and 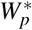 for the overtwisted DNA simulations performed in section. Panels (c) and (d): the detection of an over-the top deformation contributing one of the large jumps in the 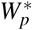 measure shown in panel (b). The end of the curve at neighboring time steps and the end plane containing the point are shown in both panels (c) and (d). A red section of curve extending vertically downward form this point s shown (an extension used in the 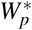 calculation in the WASP package). A section of the curve’s interior pass through this extension.

The set of writhe values for configurations 101 *−* 260 (the over-the-top unlooping of the knot) highlight the difference between *W*_*p*_ and 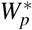. For the first part of this deformation they are identical, until the looped section passes above the end points plane (as shown in Figure 11). Then *W*_*p*_ begins to measure an changing value due to the end angle measures between this loop and the end point, 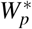 does not account for this change. Near the end of this un-looping the curve passes directly over the top of its end point an we see a jump of *−*2 in 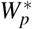, where the measure has classed the curve as jumping into a different pulled tight topological class. In the final set of deformations 260-360 the curve is pulled straight, here all three measurements are in rough agreement that there is a steady decrease in writhe, note, however, that 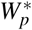 measure registers a relatively smooth change.

### Overtwisted DNA reconsidered

The discontinuous jump in the 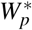 measure can lead to jumpy time series, **if** a section of the curve is continually deforming in the region directly above one of the curve’s end points. It happens that this is exactly what is occurring in the overtwisted example presented in section. The comparison of *W*_*p*_ and 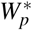 are shown in Figure 12(b). The 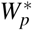 measure has some significant discontinuities, each one was analyzed an in each case it was found to result from a section of the curve passing over the bottom end of the curve, an example is shown in Figure 12(c)-(d).

A second observation is that 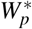 is consistently a small amount larger than *W*_*p*_. This is due to a difference between *W*_*pnl*_ and 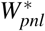 (*W*_*pl*_ is always the same for both), specifically the ignoring of the end angle depicted in Figure 11. It is interesting that the molecule appears to deform such that it partly performs the over-the-top loop, a potential means to shed topology, but never completes this deformation, instead forming second plectoneme loop.

A final mathematical point is that if one only calculated 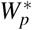 for the given curve set, then, without looking at the individual curves themselves, it is hard to tell if the jump occurs due to self crossing or over the end deformations. The *W*_*p*_ calculations, which have no changes of the order of 2 indicate it is not self-crossing but over the top deformation. Thus one can make this distinction solely from the numerical calculations (rather than visulising the curve set). Of course we should have expected this for the DNA model used as repulsive forces prevent self-crossings, but in other scenarios the combination of information provided by both the *W*_*p*_ and 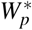 quantities could be of significant utility.

## Conclusions

Writhe is a fundamental measurement for characterizing the topology of ribbon structures, and is extensively utilized in multiple fields from biophysics, to solar physics, to aneurysm detection (to name just a few)^30–32,53,54^. However, the most commonly utilized formalism for writhe, as derived by Călugăreanu, is only applicable to the subset of problems in which the system under study is a closed loop. Although this limitation has been overcome through the development of polar writhe, it is largely underutilized in the biophysical community. Here, we have sought to rectify that problem by demonstrating the utility of polar writhe on DNA plectoneme and DNA minicircle calculations. Our results show that polar writhe is not only applicable to the analysis of open curves, such as those formed by extended DNA structures, but that it can be decomposed into local and nonlocal contributions that can provided additional information about DNA topology that can not be obtained from the Călugăreanu formalism. To aid in the adoption of polar writhe, we have developed a software package, WASP, that can be used to analyze molecular dynamics simulations of DNA. WASP is an open source software tool that is built to analyze trajectories from popular simulation packages. It is our hope that WASP will be utilized to rigorously analyze the growing field of DNA simulations in which changes of writhe are linked to biophysical phenomena.

## Supporting information

Supplement information referred to in main document

## Acknowledgements

This research was supported by the National Institute of General Medical Sciences of the National Institutes of Health (Grant 1R35GM119647). The content is solely the responsibility of the authors and does not necessarily represent the official views of the National Institutes of Health.

## Author contributions statement

Z.S. and J. W. designed the MD simulations. Z.S. and C.P. wrote the WASP code. Z.S., J.W, and C.P. analyzed the simulations, wrote, and reviewed the manuscript.

## Additional information

The source code for WASP is located in the following repository: https://github.com/WereszczynskiGroup/WASP The author(s) declare no competing interests.

