## Supplement information referred to in main document for "WASP: A software package for correctly characterizing the topological development of ribbon structures"

### ABSTRACT

Supplementary information to: WASP: A software package for correctly characterizing the topological development of ribbon structures. Here the polar writhe measure is related to existing writhe measures used in the literature.

### Ribbons and the open and closed Călugăreanu formulae

In this section we detail how the polar writhe formulation relates to the closed invariant relationship first introduced by Călugăreanu . We further relate it to the Fuller formulae which have been used in DNA supercoiling and elastic rod applications[1–6].

We begin by considering a twice differentiable space-curve $\mathbf{x}(s) \in \mathbb{R}^3$, $s \in [0,L]$, which is closed $\mathbf{x}(0) = \mathbf{x}(L)$ and parameterized by its arclength $s$. Its unit tangent vector is denoted $\mathbf{T}$. A ribbon structure is constructed using a vector field $\mathbf{V}$ normal to $\mathbf{x}$ ($\mathbf{V} \cdot \mathbf{T} = 0, \forall s$). This field may rotate around the curve $\mathbf{x}$ and defines a second curve $\mathbf{y}$, the ribbon's edge

$$\mathbf{y}(s) = \mathbf{x}(s) + \varepsilon \mathbf{V}(s), \tag{1}$$

as illustrated in Figure 1(a). It is usually assumed $\varepsilon$, the ribbon's width, is significantly smaller than the length of the curve $\varepsilon \ll L$. At first we also assume the ribbon is closed ($\mathbf{x}(0) = \mathbf{x}(L)$ and $\mathbf{y}(0) = \mathbf{y}(L)$). Note that for DNA molecules we are typically assuming the axis curve $\mathbf{x}(s)$ is the supercoiling axis of the structure, not the helical axis, *i.e.* a circle in a relaxed mini ring or a straight line for linear DNA (see Figure 1(b)).

The link-twist-writhe theorem for closed ribbon's first introduced by Călugăreanu relates three quantities

$$Lk(\mathbf{x},\mathbf{y}) = Tw(\mathbf{x},\mathbf{V}) + Wr(\mathbf{x}), \tag{2}$$

the linking number $Lk$, the writhe $Wr$, and the twist $Tw$ of the ribbon. We now introduce these quantities.

#### Linking Number, Writhe and twist of closed ribbons

The linking number $Lk$ of a ribbon in $\mathbb{R}^3$ can be defined by the Gaussian integral:

$$Lk(\mathbf{x},\mathbf{y}) = \frac{1}{4\pi} \oint_x \oint_y \mathbf{T}_\mathbf{x}(s) \times \mathbf{T}_\mathbf{y}(t) \cdot \frac{\mathbf{x}(s) - \mathbf{y}(t)}{\|\mathbf{x}(s) - \mathbf{y}(t)\|^3} \, \mathrm{d}s \, \mathrm{d}t, \tag{3}$$

where $\mathbf{T}_\mathbf{x}$ and $\mathbf{T}_\mathbf{y}$ are the unit tangent vectors of $\mathbf{x}(s)$ and $\mathbf{y}(t)$, respectively. $Lk$ is an integer valued function and most importantly a topological invariant, meaning that any continuous invertible deformation of the ribbon does not change $Lk$. Stretching the ribbon is an example of a such a deformation; cutting the ribbon is not. If the ribbon is allowed to pass through itself, the linking number changes by $\pm$ 1[7]. For ribbons the crossing of the ribbon with itself always means two crossings so there will be a change $\pm 2$ in $Lk$ if a ribbon crosses itself.

#### Linking, the Gauss map and points of view

The double integral (3) represents the degree of the Gauss map:

$$\mathscr{C}(s,t) = \frac{\mathbf{y}(s) - \mathbf{x}(t)}{\|\mathbf{y}(s) - \mathbf{x}(t)\|}, \tag{4}$$

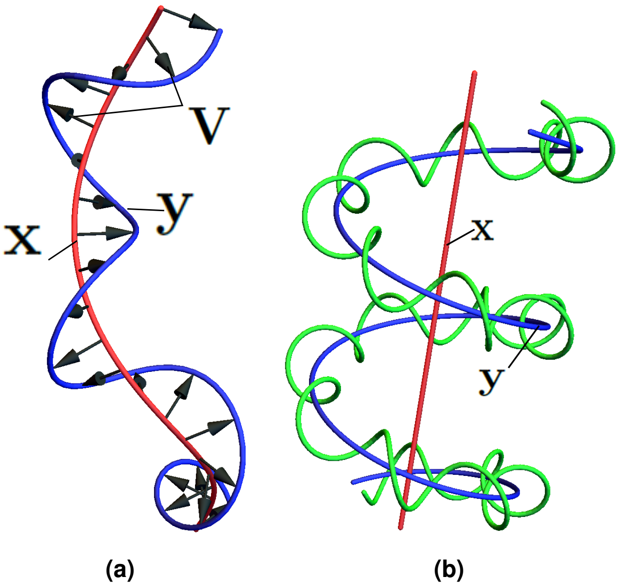


**Figure 1.** Ribbon structures. Panel (a) a ribbon is composed of an axis curve $\mathbf{x}(s)$ and a second curve $\mathbf{y}(s)$ constructed form the axis curve and a vector field $\varepsilon\vec{V}(s)$ . In panel (b) we see a ribbon structure as applied to an idealized DNA coiled-coil geometry. The axis is the straight line at the centre of the coil and the second curve $\mathbf{y}$ is the helical curve at the centre of the DNA's coiled-coil trigonometry. The green curve might represent one strand of the DNA polynucleotide chain.

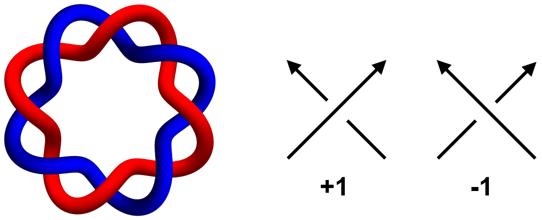


**Figure 2.** A (2,8) torus link with $Lk = Tw = 4$ and $Wr = 0$. $Lk$ can be calculated for the planar in-page projection using $Lk = \frac{\sum C_+ + C_-}{2}$ following the sign convention as pictured.

applied to all $s$ and $t$ for the two curves. The map takes pairs of points $(\mathbf{x}(s), \mathbf{y}(t))$ to points on the unit sphere $\mathbb{S}^2$, that is to say it defines directions between points (*e.g.*[8]). Equation (3) is an application of the Kroneker formula (*e.g.* Chp 8.3[9]) and each point on the unit sphere is assigned a positive or negative value by the sign of the integrand. A visual interpretation of this calculation is instructive: consider a (2,8) torus link as shown in Fig. 2. An observer may project the torus link directly into the page along some fixed direction $\mathbf{n}$ and calculate $Lk$ for the given projection by adding the number of instances in which the red and blue edge curves cross following the under/over-crossing sign convention indicated in figure 2, then dividing the sum by two. The crossings will coincide with pairs of points $(s,t)$ at which $\mathscr{C}(s,s') = \mathbf{n}$. The sign rule gives the sign of the scalar triple product $\mathbf{T}_{\mathbf{x}}(s) \times \mathbf{T}_{\mathbf{y}}(t) \cdot (\mathbf{x}(s) - \mathbf{y}(t))$. It can be shown that, whatever the direction of projection $\mathbf{n}$, the value of $Lk$ calculated by this method will be the same[9]. The integral (3) can thus be seen as the average value of this projection method over all possible choices of direction $\mathbf{n}$ (all points on the unit sphere), and is thus is a highly degenerate calculation.

#### *The net winding*

The net winding of a pair of curves $\mathbf{x}$ and $\mathbf{y}$ is defined as follows. We split $\mathbf{x}$ into $n$ sections by its turning points along $\hat{z}$ and similarly split $\mathbf{y}$ into $m$ sections by its turning points (as discussed in section 2 of the main document). For each pair $(\mathbf{x}_i, \mathbf{y}_j)$ we can calculate the mutual winding of the angle $\Theta_{ij}$ made by the vector $\mathbf{y}_j(z) - \mathbf{x}_i(z)$ **if** they share a range of $z$ values

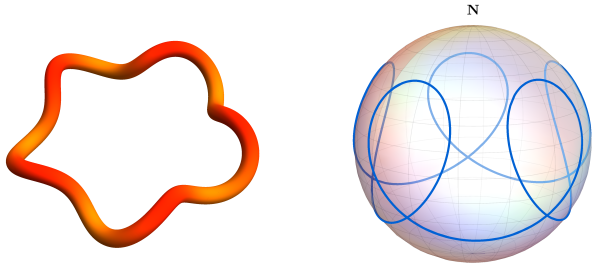


**Figure 3.** A (2,10) torus knot (Left) and its unit tangent curve **T** (Right), this is one edge of an area on the surface of the unit sphere which represents the calculation (7). Changes in shape of the curve on the left would lead to changes in this unit tangent curve, thus changing the area it bounds. This change represents the changing value of $Wr$ under deformation.

$z \in [z_{ij}^{min}, z_{ij}^{max}]$. Then the net winding is the sum of all these contributions:

$$L(\mathbf{x},\mathbf{y}) = \sum_{i=1}^{n}\sum_{j=1}^{m}\frac{\sigma_i\sigma_i}{2\pi}\int_{z_{ij}^{\min}}^{z_{ij}^{\max}}\frac{\mathrm{d}}{\mathrm{d}z}\Theta(\mathbf{x},\mathbf{y},z)\,\mathrm{d}z, \tag{5}$$

For closed ribbons this can be shown to give the same values as (3) ($L = L_k$), however, it is not the same calculation. We can define the map:

$$\mathscr{C}_z(z) = \frac{\mathbf{y}(z)-\mathbf{x}(z)}{\|\mathbf{y}(z)-\mathbf{x}(z)\|}, \tag{6}$$

which is very similar to the Gauss map $\mathscr{C}(s,t)$, except it restricts to points on the curve which have the **same** $z$ coordinate. It was shown in[10] that (5) is equal to the average of the linking calculated by projection for all directions $\mathbf{n}$ which lie in the plane *i.e.* $\mathbf{n} = (\cos(\Theta), \sin(\Theta), 0), \forall \Theta \in [0, 2\pi]$. When $\mathscr{C}_z = \mathbf{n}$ one would see a crossing of the curves. This is obviously a far less degenerate calculation than (3), but given *Lk* can be calculated from one projection, we might ask why the averaging is necessary. The answer is two-fold. First, for open ribbons, a single projection is **not** invariant to all motions which vanish at the ribbon's end (we will deal with this matter shortly) and second, using (5) yields a sensible writhing definition which is equal to *Lk*-*Tw* as we now discuss.

#### The Writhe $Wr$

The writhe $Wr$ can also be defined as a Gaussian type integral, in effect the self-linking of the curve $\mathbf{x}(s)$:

$$Wr \equiv \frac{1}{4\pi}\oint_x\oint_x \mathbf{T}_{\mathbf{x}}(s)\times\mathbf{T}_{\mathbf{x}}(t)\cdot\frac{\mathbf{x}(s)-\mathbf{x}(t)}{\|\mathbf{x}(s)-\mathbf{x}(t)\|^3}\,\mathrm{d}s\,\mathrm{d}t. \tag{7}$$

Unlike the linking number, writhe is dependent on only the axis curve of the ribbon and is not a topological invariant[8,10]. Similar to the linking number, (7) can be interpreted as the area on the unit sphere covered by the map

$$\mathscr{C}_w(s,t) = \frac{\mathbf{x}(s)-\mathbf{x}(t)}{\|\mathbf{x}(s)-\mathbf{x}(t)\|}, \tag{8}$$

with the area element positive or negative by the sign of the integrand $\vec{T}_x(s)\times\vec{T}_x(t)\cdot(\mathbf{x}(s)-\mathbf{x}(t))$. However, this area is not a degree (it is not an integer) and not a topological invariant[8,10]. In the limit $t \to s$ from below, $\mathscr{C}_w$ tends to the unit tangent vector $\mathbf{T}(s)$ and in the limit $t \to s$ from above it tends to $-\mathbf{T}(s)$. By contrast, the linking map $\mathscr{C}$ has the same limit $t \to s$ from above or below and is periodic. So the area covered by the linking integral, equation (3), will be a closed surface (and hence an integer number of signed coverings of the unit sphere), whilst the area covered the writhe integral (equation 7) has a boundary which is the union the unit tangent curves $\mathbf{T}(s)$ and $-\mathbf{T}(s)$ mapped onto the sphere (see[8]) (an example is shown in Figure 3). The $Wr$ changes continually under continuous (invertible) deformations of the ribbon as the tangent curve $\mathbf{T}$ changes, and when $x$ crosses itself it changes by a value of $\pm 2$.

Similar to *Lk*, $Wr$ can be interpreted in terms of planar projections. For example, a left-handed trefoil knot and a possible planar projection is shown in Figure 2. Walking along the projected curve and counting the self crossings with respect to their handedness using the same convention as in figure 2 gives a planar (projected) writhe of $wr = -3$. However, in contrast to the

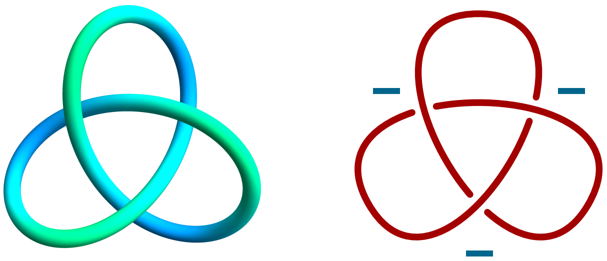

**Figure 4.** Left-handed trefoil knot with $wr = -3$ and $Wr = -3.2$. The planar writhe $wr$ can be calculated using the sign convention in Fig. 2.

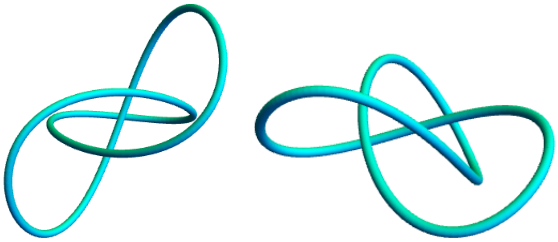

**Figure 5.** Two alternative projections/viewpoints of the trefoil knot shown in Figure 4. The left view point would lead to a projection with $wr = -3$ and the right a projection for which $wr = -4$.

linking, alternative projections **can** lead to changes in the planar writhe as indicated in Figure 5. The writhe $Wr$ is an average over ***all*** possible projections (all planar $wr$) and in the trefoil example $Wr = -3.2$, indicating the bulk of projections have $wr = -3$ for the given trefoil knot. Changing the shape of the trefoil will alter the percentage of $wr = -3$ and $wr = -4$ cases and hence change the average. We highlight that it is the averaged quantity (7) which is required for (2) to hold.

##### *Polar writhe*

The polar writhe introduced in section 2 of the main document yields exactly the same value as $W_r$ evaluated by (7) as demonstrated in[10]. Similar to the net winding, the polar writhe can be shown to be equal to the planar writhe $wr$ averaged over all projection directions $\mathbf{n}$ which lie in the plane ($\mathbf{n} = (\cos(\Theta), \sin(\Theta), 0), \forall \Theta \in [0, 2\pi]$). As indicated in Figure 4, the projected writhe $wr$ does not equal $W_r$ so the sum $wr + Tw$ would not be invariant to non self intersecting deformations of the ribbon. As demonstrated in[10], it is necessary to average over all planar directions in order to obtain the quantity $W_p = W_r$ which makes equation 2 hold. That is to say the polar writhe is the most parsimonious method for calculating $Wr$ (for closed curves). In[11] the code used to calculate $W_p$ which is used in the WASP package was shown to outperform all other existing algorithms for calculating $Wr$ and seems to generally scale linearly with the number of points in the curve.

#### The Twist $Tw$

Finally, the twist of a ribbon is equal to the total rotation of the generating vector field $\mathbf{V}$ around the tangent direction of the axis curve $x(s)$:

$$Tw \equiv \frac{1}{2\pi} \oint_{\mathbf{x}} \mathbf{T} \cdot \mathbf{V} \times \frac{\mathrm{d}\mathbf{V}}{\mathrm{d}s} \, \mathrm{d}s. \tag{9}$$

It changes continually under continuous deformations of the ribbon, even those which intersect[10].

Again considering the torus knot example of Fig. 2, because the axis curve for this torus knot is easily determined to be a planar circle, it can be seen that $Wr = 0$ for this given geometry and thus $Lk = Tw$ by (2).

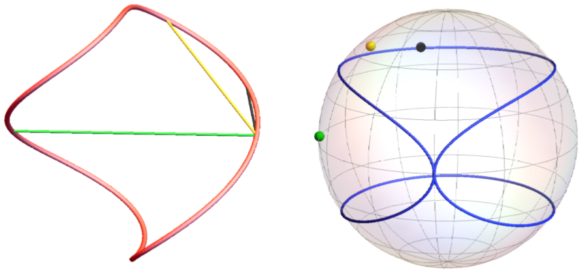

**Figure 6.** Various contributions to the $Wr$ calculation which can be classed as local or non-local. On the left a curve and a subset of its chords $C_w(s,t)$ are shown. The parameters $s$ and $t$ are chosen so these chords are vary from clearly non-local (green) to clearly local (black) and somewhere in between (yellow). On the right we see the unit tangent tangent curve $\mathbf{T}$ of the curve on the left, plotted on the unit sphere. The points corresponding to the chords shown on the left indicate their relative locality by their proximity to the unit tangent curve (the nearer to this curve the more we might say they are local).

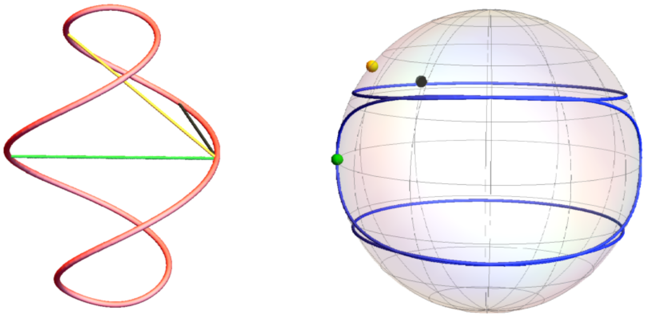

**Figure 7.** An illustration of a curve for which $W_F \neq Wr$. The curve (left) is obtained form the curve in Figure 6 by applying and extra half twist at its ends. A set of chords with the same $(s,t)$ as those in Figure 6 are shown. We note the most clearly non-local chord lies on the unit tangent curve $\mathbf{T}$ (right). This is only possible because there is a full net rotation of the curve which leads to an extra full covering of the unit sphere in the calculation of $Wr$. Such contributions are missed in the evaluation of $W_F$ (it is part of an area not bounded by $\mathbf{T}$).

### Local and non local writhing and the Fuller formulae

#### *Local and non-local crossings*

One might then reasonably call contributions to the $Wr$ calculation given by (7) from chords $C_w(s,t)$ for which $|s-t| > \varepsilon$ for some small $\varepsilon$ as *non-local*. However, as indicated in Fig 6, there are contributions which are in some sense very clearly non-local which can be used to measure the mutual entanglement of clearly distinct sections of the curve, whilst there are others which might be classified as non-local ($|s-t| > \varepsilon$) but which are determined by relatively small subsections of the curve. Put simply, a sensible value of $\varepsilon$ would be highly curve dependent, so this attempt to split the contributions into local and non-local is not wholly satisfying. As we have demonstrated in section 2 of the main document, there is a natural local/non-local separation in the polar writhe calculation which **can** provide significant extra information. For the sake of completeness we now detail (briefly) how the popular Fuller writhe expressions relate to this decomposition.

#### *The Fuller Formulae*

As mentioned above this curve $\mathbf{T}$ and its negative image $-\mathbf{T}$ are the boundaries of the area bound on the unit sphere which represents the calculation (7). Fuller[1] proposed that the total signed area $\mathscr{A}$ closed by the curve $\mathbf{T}$ on the unit sphere characterises the writhe up to an integer multiple of 2:

$$W_F = \frac{\mathscr{A}}{2\pi} - 1 \quad mod\, 2. \tag{10}$$

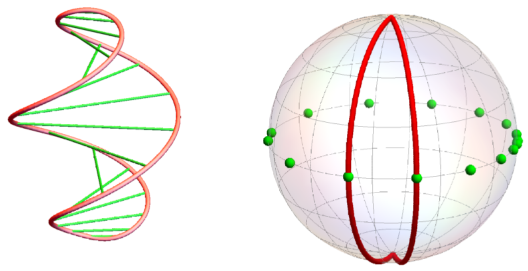

**Figure 8.** A visual interpretation of $W_{nl}$. The sets of chords $C_w(s,t)$ for which the $z$ component of $x(s)$ and $x(t)$ are equal are used to track full rotations of distinct sections of the curve separated by "turning points" for which $\mathrm{d}s/\mathrm{d}z = 0$. Such chords correspond to points on the equation and can be characterised by an angle coordinate $\Theta$ (right). The change in angle $\Delta\Theta$ between two of the pictured chords can be assigned a segmental area bound by two geodesics joining the respective angles to the poles (right). If the chord undergoes a full rotation then, as indicated, the the total area will include one full covering of the sphere.

This proposition was proven in[12]. While (10) is determined entirely by the local differential geometry of $\mathbf{x}(s)$ through the curve $\mathbf{T}$, it is not entirely local as we will shortly discuss. It will not, however, fully account for (net) signed full coverings of the sphere involved in the calculation of equation (7)[10]. For example, the Fuller formula correctly evaluates the $Wr$ of the curve shown in Figure 6, but if we apply an extra half rotation (right handed) to the curve, as indicated in Fig 7, the Fuller formula misses the correct writhe value by 1. As as a result, despite the existence of $\mathscr{O}(N)$ methods for computing writhe, $W_F$ cannot be treated as equivalent to $Wr$ as demonstrated in[13].

#### *The polar writhe and unit sphere areas*

It was shown in[10] that the non local polar writhe of a closed curve $\mathbf{x}$ with $n$ sections $\mathbf{x}_j$, split by its turning points, can be calculated as follows:

$$W_{nl}(x) = \frac{1}{2\pi}\left[\sum_{k=1}^{n-1} 2\,\sigma_i\sigma_{i+1}\,\phi_k + \sum_{\substack{i=1\\ i\neq j}}^{n}\sum_{j=1}^{n}\sigma_i\sigma_j n_{ij}\right] - 1. \tag{11}$$

where $\phi_k \in [0, 2\pi)$ is the angle made by $\mathbf{T}$ with the $x$-axis at a turning point. These contributions arise from the integrals

$$\int_{z_{ii+1}^{min}}^{z_{ii+1}^{max}} \frac{\mathrm{d}\Theta_{i(i+1)}}{\mathrm{d}z}\mathrm{d}z \tag{12}$$

which connect at the mutually shared turning point of the sections $\mathbf{x}_i$ and $\mathbf{x}_{i+1}$. The factor of $-1$ accounts for the periodicity of the closed curve as there is a contribution from the pair $\mathbf{x}_1$ and $\mathbf{x}_n$ whose mutual angle is $\pi + \phi_1$. Further it was shown (in Theorem 4 of[10]) that the $\mathscr{A}$ in the Fuller formula (10) is given by

$$\mathscr{A} = \frac{1}{2\pi}\sum_{k=1}^{n-1} 2\,\sigma_i\sigma_{i+1}\,\phi_k + W_{pl}, \quad (\mathrm{mod}\ 2) \tag{13}$$

(for closed curves). Thus we see the quantity represented by the Fuller writhe formula is the local polar withe plus a term which can be calculated from the local geometry of the curve, but actually represents a non local quantity, the turning point tangent angles $\phi_k$. For example, in the curve shown in Figure 6, the difference in the tangent angles of the upper and lower apexes of the curve give exactly $W_{pnl}$ (not $W_{pl}$). This is illustrated clearly in Figure 8 where a set of non local chords joining sections of the curve of equal $z$ height rotate in $z$. The calculation of $W_{pnl}$ for these rotating chords is equal to the area swept out by a pair geodesics joining the poles on the unit sphere bounding the areas between the angles made by these chords. The top and bottom angles are the limiting angles $\phi_k$; their difference gives $W_{pnl}$ up to an integer and so characterises part of the non local writhing.

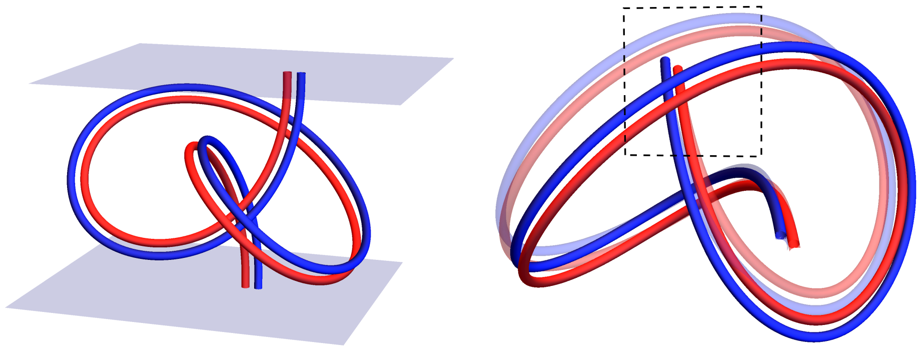

**(a)** **(b)**

**Figure 9.** Open ribbons. Panel (a): an open ended ribbon whose end points stay fixed but whose interior might deforms smoothly. Panel (b): The ribbon is observed from a particular point of view included in the evaluation of $L_k$ but not $L$. One can calculate the planar linking of this projection by assigning an orientation to the curves using the crossing rules indicated in Figure 2. The value changes in the subsequent deformation of the interior of the ribbon (shown opaquely) as we see a set of crossings is lost in the opaque figure as a section of the interior passes over the ribbon's end.

##### *The Fuller $\Delta Wr$ formula*

Fuller had a second formula for the difference in writhing between two closed curves $\mathbf{x}$ and $\mathbf{x}'$. If we can deform $\mathbf{x}$ into $\mathbf{x}'$ smoothly, such that for all the intervening curves their tangents never oppose $\mathbf{T}\cdot\mathbf{T}' \neq -1\,\forall s$ then:

$$W_r(\mathbf{x}) - W_r(\mathbf{x}') = \oint \frac{\mathbf{T}'\times\mathbf{T}}{1+\mathbf{T}\cdot\mathbf{T}'}\cdot\left(\frac{\mathrm{d}\mathbf{T}}{\mathrm{d}s}+\frac{\mathrm{d}\mathbf{T}'}{\mathrm{d}s}\right)\mathrm{d}s. \tag{14}$$

A rigorous proof for this was given in[12]. The so called non opposition condition $\mathbf{T}\cdot\mathbf{T}' \neq -1$ means the two curves cannot differ by a looped section (*i.e.* plectonemes). Since $W_p = W_r$, (14) can also be applied to the polar writhe. It was shown in[10] that the difference $W_p(\mathbf{x}) - W_p(\mathbf{x}')$ can be characterised by the difference in local writhing $W_{pl}$ and the tangent angels $\phi_k$. Specifically if we write

$$W_p^{\dagger}(\mathbf{x}) = W_{pl}(\mathbf{x}) + \frac{1}{2\pi}\sum_{k=1}^{n-1} 2\,\sigma_i\sigma_{i+1}\,\phi_k. \tag{15}$$

Then

$$\begin{aligned} W_p(\mathbf{x}) - W_p(\mathbf{x}') &= W_p^{\dagger}(\mathbf{x}) - W_r^{\dagger}(\mathbf{x}') \\ &= \oint \frac{\mathbf{T}'\times\mathbf{T}}{1+\mathbf{T}\cdot\mathbf{T}'}\cdot\left(\frac{\mathrm{d}\mathbf{T}}{\mathrm{d}s}+\frac{\mathrm{d}\mathbf{T}'}{\mathrm{d}s}\right)\mathrm{d}s. \end{aligned}$$

So we see the second Fuller formula coveys the same mixture of local and non local information as the area formula, but **not** the full local non-local decomposition of the polar writhe.

### Link twist and writhe of open ribbons

In this section we consider the topology of ribbons bound between two planes whose end pairs $(x(0),y(0))$ and $(x(L),y(L))$ lie in the bounding planes (as in Figure 9). If its ends are prevented from rotating and it deforms without crossing itself we should expect the mutual linkage of the two ends of the ribbon to be conserved. As discussed in the introduction and proven in[10], the net winding given by (5) (which is equally applicable to open ribbons) **is** invariant if the ends of the ribbon are prevented from rotating. Further it is shown that

$$L(\mathbf{x},\mathbf{y}) = W_p(\mathbf{x}) + Tw(\mathbf{x},\mathbf{V}). \tag{16}$$

##### *Gaussian linking/writhing?*

Another suggestion in the literature has been to use $Wr$, defined by (7), which can be applied to open curves. This is why calculations of this quantity were compared to polar writhe calculations on the DNA models in the main paper. There are two significant drawbacks to this approach. First, there is no known proof that the sum $W_r(\mathbf{x}) + Tw(\mathbf{x},\mathbf{V})$ is equal to $L_k(\mathbf{x},\mathbf{y})$

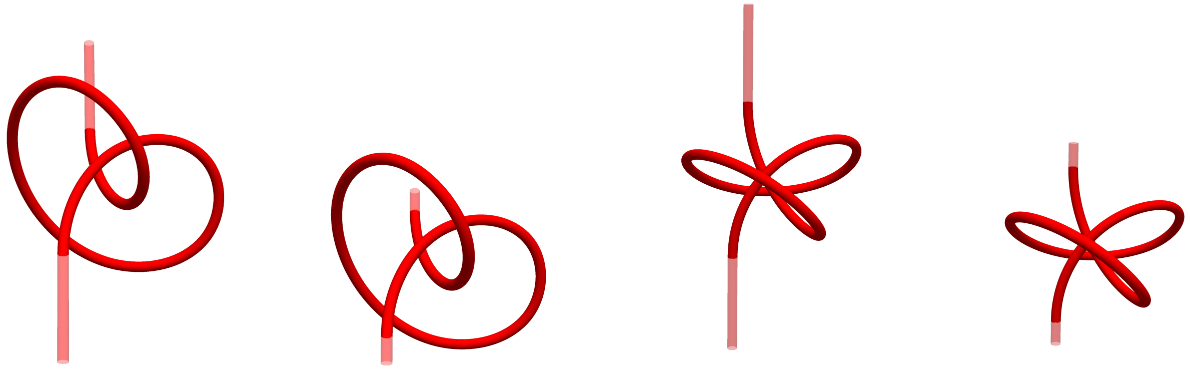


**Figure 10.** Points of view of a pair of curves from the set whose writhe values are shown in Figure **??**. The opaque sections of curve are the extensions used to create a pulled tight know as shown in Figure 10(a) of the main paper. Panels (a) and (b) indicate the projection from which a planar writhe calculation *wr* could be made which would be part of the average involved in $Wr$, equation (7), but **not** $W_p$. As the knot is "loosened" (the extension shrunk), a crossing present in panel (a) disappears in (b). Panels (c) and (d) are the same curves shown in (a) and (b) respectively, but in this case the projection from which a planar writhe calculation *wr* could be made would be part of the average involved both $W_r$ **and** $W_p$. There is no loss/gain in crossings.

defined by (3) (which again is applicable to open ribbons). Second, even if that were the case, the linking $L_k(\mathbf{x},\mathbf{y})$ is **not** an invariant to motions which change the shape of the interior of the ribbon, even if the ends are fixed. To see why this is, we note that the integral (3) still represents the signed area covered on the unit sphere by the map $\mathscr{C}(s,t)$. However, this is no-longer an integer and has a boundary represented by all the chords $\mathscr{C}(0,t)\ \forall t\in[0,L_2]$, $\mathscr{C}(L_1,t)\ \forall t\in[0,L_2]$, $\mathscr{C}(s,0)\ \forall s\in[0,L_1]$ and $\mathscr{C}(s,L_2)\ \forall s\in[0,L_1]$. Since this boundary depends on the geometry of the interior of the curves $\mathbf{x}$ and $\mathbf{y}$ it can change even if the boundary points stay fixed. Another way of putting this is in terms of link projections. As in the closed ribbon case, if we project the link along a direction $\mathbf{n}$ and count crossings we can define a planar linking; $L_k(\mathbf{x},\mathbf{y})$ is again the average over all such projections. However, now, the planar linking will not be the same for all directions and projections can lose crossings if the ribbon's interior deforms to pass under/above one of its end points in projection, as indicated in Figure 9(b).

Since the net-winding only measures the crossing of chords $C_z(z)$, the only contributions involving the end points occur **between** the end points, *i.e.* the angles $\Theta(x(0),y(0))$ and $\Theta(x(L),y(L))$ which lie in the planes shown in Figure 9(a). Thus if they are fixed, there is no change to the boundary of the calculation (see Theorem 4 of[10]). Put simpler: the projection average involved in the net winding will not include the possibility of the interior crossing the ribbon's end, as is the case in in 9(b).

##### *Knot disentanglement revisited*

We recall the knot disentanglement deformation depicted in Figure 10 of the main paper and whose writhe calculations are shown in Figure 12 of the main paper. The first tranche of the knot's deformation covers the loosening of the knot. This is achieved by extending the end of the knot with a straight line as shown in Figure 10 (a procedure which produces the transition form Figure 10 (a) to (b) of the main text). Since the writhe is invariant under a uniform scaling, this is equivalent to pulling the end of the knot tight (in reverse in the calculations). Thus the shape of the knotted section is unaltered. The calculations of the $Wr$ ( defined by (7)) for this curve show a steady change in value where the $W_p$ calculations do not. This occurs because the $Wr$ calculation involves an average over directions which see the development of additional crossings of the curves interior and its end point, as shown in Figure 10(a) and (b). By contrast the $W_p$ calculation is an average over projection directions which do not see these extra crossings, see Figures 10(c) and (d).

#### Closures

Approaches such as those outlined by[2,11,13–15] discuss methods of artificial closure of the ribbon such that an invariant link measure is recovered and the Călugăreanu theorem is valid. Work by Starostin[16] discusses extending writhe to open curves through closures of the unit tangent curve on the surface of the unit sphere. It was shown in[10] that it is always possible to extend an open curve $\mathbf{x}$ to with a sections of curve $\mathbf{x}_c$ to form a closed curve $\mathbf{x}\cup\mathbf{x}_c$ such that $W_p(\mathbf{x}) = W_r(\mathbf{x}\cup\mathbf{x}_c)$ and in[11] that it is always possible to extend a ribbon in a similar manner such that $L(\mathbf{x},\mathbf{y}) = Lk(\mathbf{x}\cup\mathbf{x}_c,\mathbf{y}\cup\mathbf{y}_c)$. From the point of view of topologically constraining the ribbon structure, the development of a closure is an unnecessary complication. In addition, as we have highlighted, the polar writhe-net winding formulation of this constraint has the additional benefit of a clear local/non-local decomposition.

#### The Fuller writhe of open curves

Several authors suggested the following expression for quantifying the writhing of the an open curve:

$$W_F(\mathbf{x}) = \int_0^L \frac{\hat{z}\cdot\mathbf{T}\times\frac{\mathrm{d}\mathbf{T}}{\mathrm{d}s}}{1+\hat{z}\cdot\mathbf{T}}\mathrm{d}s. \tag{17}$$

This is nearly identical to the expression for $W_{pl}$ without the absolute argument in the denominator ($\hat{z}\cdot\mathbf{T}$ not $|\hat{z}\cdot\mathbf{T}|$). The quantity in (17) is labelled $W_F$ here as it can be derived from the Fuller $\Delta W_r$ expression (14)[2], but it was also independently derived in[3] and[4] as a limited means of topologically constraining the molecule using descriptions of the curve $\mathbf{x}$ in terms of the three-dimensional group of rotations. However, as demonstrated in[13], it cannot be used to fully topologically constrain the structure (in particular in plectomemed configurations). This can be made clear in its relationship to the polar writhe. It was shown in[10] that $W_F(\mathbf{x})$, given by (17) is equal to $W_p^{\dagger}(\mathbf{x})$ given by (15). This makes clear that the missing contribution is the integer windings of each pair of distinct sections of the curve $\mathbf{x}_i$, and thus with the WASP package it would be possible to precisely quantify the error in the expression (17).
